# Proton FLASH radiotherapy enhances control of triple-negative breast cancer through STING–IRF3 and CD8+ T-cell immunity

**DOI:** 10.64898/2026.09.19.751623

**Authors:** Pierre Loap, Ioannis Paraskevaidis, Cezara Cheptea, Kristianna Kolker, Michele Kim, Jufri Setianegara, Kriti Singh, Iulia Maria Berianu, James Metz, Roger A. Greenberg, Warren B. Bilker, Eva Berlin, Charles-Antoine Assenmacher, Constantinos Koumenis, Eric Diffenderfer, Neil K. Taunk, Ioannis I. Verginadis

## Abstract

FLASH radiotherapy delivers radiation at ultra-high dose rates and has been demonstrated to spare normal tissue compared to standard radiotherapy, but it is not known if dose rate also modifies tumor response. Here we compare a single 13.5 Gy fraction of proton irradiation delivered at FLASH (F-PRT) or Standard (S-PRT) dose rate in immunocompetent C57BL/6 mice bearing EO771 or AT3 triple-negative mammary tumors. At this identical physical dose, F-PRT delays tumor growth more than S-PRT at both heterotopic and orthotopic sites. The effect is largest in EO771, where F-PRT also prolongs tumor-volume endpoint-free survival and reduces the emergence of lung metastases relative to S-PRT. F-PRT induces earlier intratumoral STING expression and IRF3 nuclear translocation, higher type I interferon levels and greater CD8+ T-cell infiltration. CD8+ T-cell depletion or systemic STING inhibition abolishes the F-PRT advantage. Combined with anti-PD-1 and agonistic anti-CD40, both modalities produce durable complete responses that reject contralateral rechallenge, but F-PRT limits tumor progression before response and accelerates regression. FLASH proton radiotherapy not only improves normal-tissue tolerance, but also antitumor immunity, suggesting that ultra-high dose rate could widen the therapeutic window from both sides.

## Introduction

Adjuvant radiotherapy reduces locoregional recurrence and potentially breast cancer-specific mortality after both breast-conserving surgery and mastectomy [1,2] by targeting residual microscopic disease remaining in the breast and regional nodes after surgery [3]. This is particularly relevant in triple-negative breast cancer (TNBC), an aggressive subtype defined by the absence of hormone-receptor and HER2 expression and associated with high rates of early locoregional and distant recurrence [4–7]. Although pathological complete response after neoadjuvant therapy predicts excellent long-term outcome, patients with residual invasive disease remain at high risk of relapse despite escalation with capecitabine, olaparib or immune checkpoint blockade [8–11]. Further improvement in the antitumor efficacy of locoregional radiotherapy is therefore needed in this population.

TNBC is among the most immunogenic breast cancer subtypes: tumors are frequently infiltrated by lymphocytes, and higher levels of stromal tumor-infiltrating lymphocytes are associated with response to systemic therapy and with recurrence-free and overall survival, including in patients who receive no adjuvant chemotherapy [12–16]. In high-risk early TNBC, pembrolizumab added to neoadjuvant chemotherapy and continued after surgery improves pathological complete response, event-free survival and overall survival [17–19]; in advanced PD-L1-positive disease, checkpoint blockade likewise improves outcomes [20,21]. Radiotherapy also modulates antitumor immunity: it releases tumor antigens, induces type I interferon signaling and can act as an in situ vaccine, although it also recruits immunosuppressive populations and depletes circulating lymphocytes [22–24]. Radiotherapy approaches that increase immune cell recruitment or reduce local immunosuppression may therefore improve tumor control in TNBC [25,26].

The dose of radiotherapy that can be delivered, in general, is limited principally by toxicity to the surrounding organs at risk [27]. FLASH radiotherapy delivers radiation at ultra-high dose rates, generally defined as a mean dose rate of at least 40 Gy/s (compared to standard rates of 0.5 Gy/s), and is being developed as a means of improving the therapeutic ratio of radiotherapy [28,29]. FLASH was initially investigated for its ability to reduce normal-tissue toxicity while maintaining tumor control in preclinical models [30–41], an effect reproduced with electrons, photons and protons across lung, brain, intestine, skin, salivary gland and heart, and extended to veterinary patients and to a first human treatment [42,43]. The underlying mechanisms remain incompletely defined; transient oxygen depletion, peroxyl-radical recombination and differential sparing of circulating immune cells have all been proposed [28,44–46]. Whether ultra-high dose rate also changes the antitumor response at an identical physical dose is far less well established. Recent preclinical studies suggest that proton FLASH induces a more favorable immune response than standard-dose-rate irradiation. In lung tumor models, proton FLASH increased intratumoral CD8+ T-cell infiltration and reduced regulatory T cells and immunosuppressive myeloid populations compared with standard-dose-rate irradiation [47]. In brain tumor models, it promoted greater T-cell infiltration and proinflammatory macrophage polarization while limiting immunosuppressive macrophage states [48]. Abdominopelvic FLASH irradiation likewise improved the efficacy of PD-1 blockade in preclinical ovarian cancer models [49]. These findings suggest that proton FLASH may enhance antitumor immunity beyond that achieved with standard-dose-rate irradiation, although not every study has detected differences in immune infiltration or tumor control between dose rates. Critically, the effect of dose rate on innate immune sensing within the tumor itself, as distinct from the composition of the infiltrate that follows, has not been directly examined.

Whether these immune effects translate into improved tumor control in TNBC remains unknown. Here, we compared proton FLASH with standard-dose-rate proton irradiation at an identical physical dose of 13.5 Gy and investigated the immune mechanisms underlying differences in antitumor response. Proton FLASH produced greater tumor control than standard-dose-rate irradiation in heterotopic and orthotopic TNBC models (EO771 and AT3), most consistently in EO771. This differential effect was no longer observed after CD8+ T-cell depletion or pharmacological STING inhibition and was accompanied by earlier STING–IRF3 activation, increased type I interferon production and greater intratumoral CD8+ T-cell accumulation. When combined with anti-PD-1 and agonistic anti-CD40, proton FLASH accelerated tumor regression relative to standard irradiation. Together, these data identify dose rate as a determinant of radiation-induced antitumor immunity, independent of the delivered dose.

## Methods

### Animals and ethical approval

Female C57BL/6J mice aged 9–11 weeks were obtained from The Jackson Laboratory and housed in an AAALAC International-accredited facility at the University of Pennsylvania, with food and water available ad libitum. Mice were acclimated for at least 1 week before experimentation. All animal procedures were approved by the University of Pennsylvania Institutional Animal Care and Use Committee (protocol no. 805191 and 807852) and performed in accordance with institutional guidelines. Group sizes for each experiment are reported in the corresponding figure legends. Mice were randomly allocated to treatment groups on the day of irradiation. Animal experiments are reported in line with the ARRIVE 2.0 essential requirements.

### Cell lines and tumor implantation

The syngeneic murine mammary carcinoma cell lines EO771 (ATCC, CRL-3461) and AT3 (Sigma-Aldrich, SCC178), both of C57BL/6 origin and widely used in immunocompetent models of triple-negative breast cancer, were used in this study [50–52]. Cells were cultured in Dulbecco’s modified Eagle medium (Corning, 10-013-CV) containing 4.5 g l−1 glucose, L-glutamine and sodium pyruvate and supplemented with 10% fetal bovine serum and 1% penicillin-streptomycin. Cells were harvested during exponential growth and resuspended in sterile phosphate-buffered saline (PBS) immediately before implantation. For heterotopic tumor implantation, 5 × 10^5^ cells in 200 µl PBS were injected subcutaneously into the right flank. For orthotopic tumor implantation, 2 × 10^5^ cells in 50 µl PBS were injected into the left second mammary fat pad (thoracic). Implantation was performed under isoflurane anesthesia (3% in oxygen at 1 l/min) after shaving and skin preparation. Tumors were irradiated approximately 10 days after implantation, when they reached a volume of approximately 80 mm³. Cells were used within 3-5 passages after thawing and tested negative for Mycoplasma.

### Proton irradiation and dosimetry

Proton irradiation was delivered in single-energy transmission mode using the fixed horizontal beamline of an IBA Proteus Plus C230 cyclotron (230 MeV; range, approximately 32 g/cm^2^) in a dedicated preclinical irradiation room [36,40]. Mice were anesthetized with isoflurane (3% in oxygen at 1 l/min) and immobilized on a custom vertical support. Tumors were positioned at the center of an 8-mm-diameter circular field using a collimator placed approximately 1 cm from the tumor surface. Flank tumors were aligned perpendicular to the proton beam. For mammary fat pad tumors, mice were tilted by approximately 45° to align the tumor with the beam axis. Tumors received a single physical dose of 13.5 Gy at either standard dose rate (S-PRT) or FLASH dose rate (F-PRT). This dose was selected to maintain continuity with our previous proton FLASH studies, in which 13.5 Gy per fraction was used in a 3 × 13.5-Gy cardiac irradiation model and subsequently in studies of FLASH-mediated sparing of circulating lymphocytes [38,53]. It also provides a clinically interpretable reference for breast radiotherapy; assuming an α/β ratio of 5 Gy, 13.5 Gy in one fraction corresponds to a biologically effective dose of 50.0 Gy₅, close to the 53.0 Gy₅ delivered by the FAST-Forward regimen of 26 Gy in five fractions [54]. Non-irradiated mice underwent the same anesthesia and positioning procedures without beam delivery. Field alignment, size and flatness were verified before each irradiation using Gafchromic EBT3 film. Absolute dosimetry was performed using a NIST-traceable Advanced Markus ionization chamber according to the IAEA TRS-398 protocol. Beam output was monitored online using a cross-calibrated PTW Bragg Peak transmission chamber. Mean dose rates were 114 ± 4.6 Gy/s for F-PRT and 0.52 ± 0.14 Gy/s for S-PRT. Beam current was the only delivery parameter varied between S-PRT and F-PRT. The mean measured dose was 13.33 ± 1.31 Gy and 13.28 ± 1.30 Gy for F-PRT and S-PRT, respectively.

### Tumor monitoring and growth analysis

Tumor dimensions were measured three times per week using digital calipers. Tumor volume was calculated as V = d² × D/2, where d and D represent the shortest and longest orthogonal diameters, respectively. Mice were monitored until tumor volume reached 1,000 mm³ or until a humane endpoint (tumor ulceration, body-condition score below 2, or weight loss exceeding 20%) was reached, whichever occurred first. Tumor-volume endpoint-free survival was defined as the interval from injection to a tumor volume of 1,000 mm³. Complete response was defined as the absence of a visible or palpable tumor at clinical examination, irrespective of its subsequent duration. A durable complete response was defined separately as complete response without subsequent tumor regrowth during follow-up. Mice were euthanized by CO₂ inhalation followed by cervical dislocation. Tumor doubling time was estimated by fitting an exponential growth model to measurements obtained during the exponential growth phase. The volume window used for the fit is stated in each figure legend and was applied identically to all groups within a given experiment. Complete responders and tumors without a measurable exponential growth phase were excluded only from the tumor doubling-time analysis.

### CD8+ T-cell depletion and STING inhibition

For CD8+ T-cell depletion, EO771-bearing mice received intraperitoneal injections of anti-CD8 antibody (clone 2.43, Bio X Cell, #BE0061, Lot:94122401; 200 µg per mouse) or rat IgG2b isotype control (clone LTF-2, Bio X Cell, #BE0090, Lot:915224J3; 200 µg per mouse). Treatment was initiated 1 day before irradiation and repeated every 4 days until endpoint. Mice were assigned to non-irradiated, S-PRT or F-PRT groups. CD8+ T-cell depletion was confirmed by flow cytometry of peripheral blood collected by retro-orbital bleeding under isoflurane anesthesia on the day of the second antibody injection. Following red blood cell lysis, leukocytes were stained with antibodies against CD45, CD3, and CD8 and analyzed by flow cytometry. For STING inhibition, EO771-bearing mice received H-151 (10 mg kg−1; MedChemExpress, HY-112693, Lot: 1035356) or vehicle by intraperitoneal injections once daily, beginning 1 day before irradiation and continuing until endpoint. H-151 was formulated in 10% DMSO, 40% PEG300, 5% Tween-80 and 45% saline, added sequentially in that order, and administered intraperitoneally at a volume of 100 µl per mouse.

### Immunotherapy and tumor rechallenge

EO771-bearing mice received anti-PD-1 antibody (clone RMP1-14, Bio X Cell, BE0146, Lot: 924625A1 and 949225A2B; 200 µg per mouse) by intraperitoneal injection three times per week for six doses, beginning 24 h after irradiation. Agonistic anti-CD40 antibody (clone FGK4.5, Bio X Cell, BE0016-2, Lot: 903625J3, 100 µg per mouse) was administered once by intraperitoneal injection 24 h after irradiation. Control mice received rat IgG2b control antibody (clone LTF-2, Bio X Cell, catalog no. BE0090, lot no. 915224J3; 200 µg per mouse) according to the corresponding administration schedule. Maximum tumor volume before regression and time to regression below 50 mm³, below 30 mm³ and to complete disappearance were recorded. Mice remaining tumor-free for 4 weeks after complete tumor disappearance were rechallenged with 2 × 10^5^ EO771 cells in 50 µl PBS injected into the contralateral right second mammary fat pad. Tumor growth was monitored three times per week for 2 weeks after rechallenge. In the absence of tumor regrowth, spleens and tumor-draining lymph nodes were collected 2 weeks after rechallenge for flow cytometric analysis of CD4+ and CD8+ central-memory and effector-memory T-cell populations.

### Tissue processing and flow cytometry

Tumors and spleens were collected 7 days after irradiation for immune profiling. Tumors were minced on ice and digested in serum-free DMEM containing collagenase II (5 mg/ml; Worthington Biochemical, LS004176, Lot: 45J25422), collagenase IV (5 mg/ml; Gibco, 17104019, Lot: 3298441) and DNase I (2 mg/ml; Roche, 10104159001, Lot:73953000) for 45-60 min at 37 °C with agitation. Spleens were mechanically dissociated. Cell suspensions were filtered through 70-µm and 45-µm strainers, washed and resuspended in PBS containing 4% fetal bovine serum at approximately 2 × 10^6 cells/ml. Cell viability was assessed using LIVE/DEAD Fixable Aqua stain (Thermo Fisher Scientific, L34965, Lot: 3417064). Cells were stained for surface markers and fixed and permeabilized for intracellular staining where indicated. T-cell panels included antibodies against CD45, CD3, CD4, CD8 and CD25, together with intracellular staining for FOXP3, IFN-γ and granzyme B. Myeloid panels included antibodies against CD45, CD11b, F4/80, CD80, CD206, arginase 1, Ly6G, Ly6C, Gr-1 and CD11c. Samples were acquired using a BD FACSCanto II flow cytometer and analyzed using FlowJo software (version 10). Gating was performed sequentially on singlets, viable cells and CD45+ leukocytes. Complete gating strategies for all flow-cytometry panels are shown in **Supplementary Fig. 6**. Target, fluorochrome, supplier, catalog number, lot number and clone for every antibody used in the flow cytometry experiments are provided in **Supplementary Table 1**.

### Immunofluorescence

EO771 tumors were collected 2 or 7 days after irradiation, embedded in OCT and sectioned at 10 µm. Sections were stained for CD8, STING or phosphorylated IRF3 and counterstained with Hoechst. Tissue sections were fixed using antigen-specific conditions. Sections designated for CD8 analysis were fixed with 100% methanol, whereas sections used for STING analysis were fixed with 2% paraformaldehyde, and those for pIRF3 staining were fixed with 4% paraformaldehyde. Following fixation, sections were blocked with 8% bovine serum albumin (BSA) in PBS-TT (0.1% Tween-20, 0.025% Triton X-100) at room temperature for 2 hours. Sections were incubated with primary antibodies against CD8 (1:100; Abcam, ab22378, Lot: 1074953-1), STING (1:200; Proteintech, 19851-1-AP, Lot: 00205955) or phosphorylated IRF3 (1:200; Proteintech, 80519-2-RR, Lot: 00148186) and then with the corresponding fluorescent secondary antibodies.

Images were acquired at ×20 magnification using a Zeiss Observer.Z1 inverted microscope with identical acquisition settings for all groups within each experiment. All quantification was performed in ImageJ/FIJI on the raw, unprocessed .czi files [55,56]. Brightness and contrast were adjusted only for the preparation of display figures, applied identically to all images. CD8+ infiltration was quantified as the number of positive cells per ×20 field. STING and phosphorylated IRF3 expression were quantified using ImageJ Fiji by assessing the proportion of positive cells and immunofluorescent signal intensity. Nuclear phosphorylated IRF3 was quantified by colocalization with Hoechst through ImageJ/Fiji JaCop plugin [57]. For each tumor, 10–15 fields spatially distributed across the tumor section were acquired to provide representative tumor coverage. Field-level measurements were averaged to obtain a single value per mouse, which was used as the biological replicate for statistical analysis.

### Tumor cytokine and chemokine analysis

Tumors were collected 7 days after irradiation and homogenized in 1× RIPA buffer (Thermo Scientific, catalog no. 89900) supplemented with protease inhibitor cocktail (Roche, catalog no. 11836170001). Lysates were clarified by centrifugation, and total protein concentrations were determined using the Pierce BCA Protein Assay Kit (Thermo Scientific, catalog no. 23227). IFN-α, IFN-β, IFN-γ, CXCL9/MIG, and CCL5/RANTES concentrations were measured using a custom 7-plex MILLIPLEX Mouse Cytokine Expansion Panel 1 magnetic bead assay (Millipore, catalog no. MCYT1-190K-07, kit identifier mcyt1907, lot no. 89) according to the manufacturer’s instructions. Data were acquired using a MAGPIX system and analyzed with xPONENT software. Analyte concentrations were calculated from five-parameter logistic standard curves, normalized to total protein concentration, and expressed as fold change relative to the corresponding non-irradiated group.

### H&E staining and pulmonary metastasis assessment

Spontaneous pulmonary metastases arising from orthotopic mammary tumors were assessed without experimental induction of lung metastases by tail-vein or other intravenous tumor-cell injection. Lungs were collected when the primary orthotopic tumors reached the predefined 1,000 mm³ endpoint. Macroscopic surface nodules were counted, and metastatic burden was additionally quantified on histological sections as the percentage of lung parenchymal area occupied by metastases. The lungs were processed into paraffin blocks, sectioned at 5 μm and routinely stained with hematoxylin and eosin. Slides were then digitized using the Leica Versa 200 whole slide scanner (Leica BioSystems), and the Aperio ImageScope software (Leica BioSystems) was used for image visualization and annotations. For each mouse, three sections separated by 100 µm were analyzed and the mean value across sections was used.

### Statistical analysis

Statistical analyses were conducted using GraphPad Prism (version 10). Changes in tumor volume over time were assessed using mixed-effects models with treatment group, time and their interaction as fixed effects. Tumor-volume endpoint-free survival was analyzed using the Kaplan-Meier method, and survival curves were compared by log-rank tests with Holm-Šidák adjustment for multiple comparisons. Differences among three or more groups were evaluated by one-way analysis of variance followed by Tukey’s multiple-comparison test when data were normally distributed, and by the Kruskal-Wallis test with Dunn’s correction otherwise; normality was assessed using the Shapiro-Wilk test. Two-group comparisons were performed using two-tailed unpaired Student’s t-tests or Mann-Whitney tests, as indicated in the corresponding figure legends. For immunofluorescence analyses, the mouse was considered the biological replicate; measurements from multiple microscopy fields within each tumor were averaged before statistical testing. Other statistical procedures are described in the relevant figure legends. All analyses were two-sided, and P values below 0.05 were considered statistically significant. The P values and the number of biological replicates are reported in the figures or figure legends. No formal power calculation was performed. Animals were randomly allocated to experimental groups. Investigators could not be blinded during irradiation, since group allocation was apparent at the time of beam delivery; all subsequent outcome assessment — caliper measurement, immunofluorescence acquisition and quantification — was performed blinded to treatment allocation. No animals or measurements were excluded from the primary longitudinal or survival analyses. Complete responders and tumors that remained measurable but showed no sustained progressive growth and therefore never entered a measurable exponential growth phase were excluded only from tumor doubling-time analyses, as stated in the corresponding figure legends.

## Results

### FLASH proton irradiation improves tumor control in heterotopic TNBC models

We first compared the antitumor effects of FLASH and standard-dose-rate proton irradiation using two eterotopic TNBC models in immunocompetent mice with differing sensitivity to immune checkpoint blockade; EO771 has been reported to be more responsive than AT3 to anti-PD-1 therapy [58]. Subcutaneous EO771 and AT3 tumors received a single 13.5-Gy fraction delivered at either standard dose rate (0.52 ± 0.14 Gy/s; S-PRT) or FLASH dose rate (114 ± 4.6 Gy/s; F-PRT), while non-irradiated tumors served as controls (NR; **Fig. 1a**). In EO771 tumors, F-PRT produced a greater tumor growth delay than S-PRT and NR (**Fig. 1b**). F-PRT also improved tumor-volume endpoint-free survival compared with S-PRT and with NR (**Fig. 1c**). Complete tumor regression occurred in 2 of 10 F-PRT-treated mice and in none of the 9 S-PRT-treated mice (**Supplementary Fig. 1a**). To assess tumor growth kinetics after the initial treatment response, doubling time was estimated over the 200-1,000 mm³ growth window. Two F-PRT-treated tumors that did not reach this window were excluded from this analysis. F-PRT prolonged tumor doubling time compared with NR, whereas S-PRT did not differ from NR (**Fig. 1d**).

**Fig. 1.**
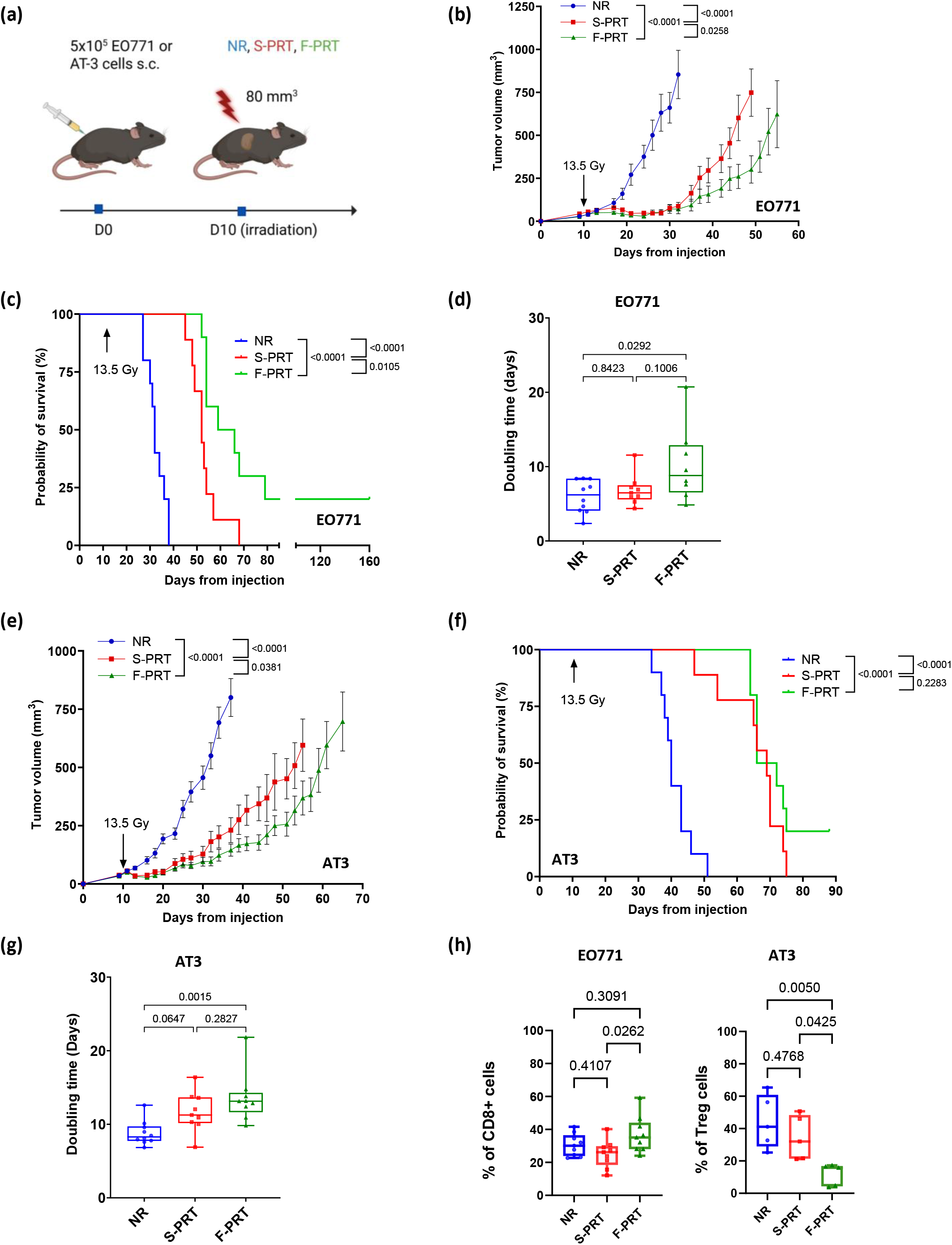
FLASH proton therapy enhances antitumor immunity and tumor control compared with standard dose-rate proton therapy in heterotopic TNBC models. (a) Experimental workflow. EO771 or AT3 triple-negative mammary carcinoma cells (5 × 10⁵) were injected subcutaneously into the flank of immunocompetent C57BL/6 mice. Ten days after implantation, when tumors reached approximately 80 mm³, mice were assigned to the non-irradiated control (NR), standard dose-rate proton irradiation (S-PRT), or FLASH proton irradiation (F-PRT) groups. Irradiated tumors received a single 13.5-Gy fraction of proton irradiation. (b) Mean tumor growth kinetics of EO771 tumors in the NR (n = 10), S-PRT (n = 9), and F-PRT (n = 10) groups. Data are shown as mean ± SEM. NR, S-PRT, and F-PRT are shown in blue, red, and green, respectively. Tumor growth curves were compared using mixed-effects models. (c) Kaplan-Meier analysis of tumor endpoint-free survival in mice bearing EO771 tumors. Mice were followed until tumors exceeded the predefined endpoint volume of 1,000 mm³, at which point they were euthanized. NR (n = 10), S-PRT (n = 9), and F-PRT (n = 10) are shown in blue, red, and green, respectively. Survival curves were compared using log-rank tests with Holm-Šídák correction for multiple comparisons. (d) Tumor doubling time of EO771 tumors in the NR, S-PRT, and F-PRT groups. Doubling time was calculated by fitting an exponential growth model to tumor volume measurements between 200 and 1,000 mm³. Boxes show the median and interquartile range, whiskers indicate minimum and maximum values, and each point represents one mouse. Sample sizes were n = 10 for NR, n = 9 for S-PRT, and n = 8 for F-PRT. Two F-PRT-treated mice were excluded from doubling-time calculation because they did not reach a sufficient tumor volume for exponential fitting. Comparisons were performed using ordinary one-way ANOVA with Tukey’s correction for multiple comparisons. (e) Mean tumor growth kinetics of AT3 tumors in the NR (n = 10), S-PRT (n = 9), and F-PRT (n = 10) groups. Data are shown as mean ± SEM. NR, S-PRT, and F-PRT are shown in blue, red, and green, respectively. Tumor growth curves were compared using mixed-effects models. (f) Kaplan-Meier analysis of tumor endpoint-free survival in mice bearing AT3 tumors. Mice were followed until tumors exceeded the predefined endpoint volume of 1,000 mm³, at which point they were euthanized. NR (n = 10), S-PRT (n = 9), and F-PRT (n = 10) are shown in blue, red, and green, respectively. Survival curves were compared using log-rank tests with Holm-Šídák correction for multiple comparisons. (g) Tumor doubling time of AT3 tumors in the NR, S-PRT, and F-PRT groups. Doubling time was calculated by fitting an exponential growth model to tumor volume measurements between 200 and 1,000 mm³. Boxes show the median and interquartile range, whiskers indicate minimum and maximum values, and each point represents one mouse. Sample sizes were n = 10 for NR, n = 9 for S-PRT, and n = 9 for F-PRT. One F-PRT-treated mouse was excluded from doubling-time calculation because the tumor did not reach a sufficient volume for exponential fitting. Comparisons were performed using ordinary one-way ANOVA with Tukey’s correction for multiple comparisons. (h) Flow cytometry quantification of CD8⁺ T cells and regulatory CD4⁺ T cells within the tumor microenvironment of EO771 and AT3 tumors 7 days after irradiation. Boxes show the median and interquartile range, whiskers indicate minimum and maximum values, and each point represents one mouse. For EO771 CD8⁺ T-cell quantification, sample sizes were n = 9 per group. For AT3 regulatory CD4⁺ T-cell quantification, sample sizes were n = 5 per group. Comparisons were performed using ordinary one-way ANOVA with Tukey’s correction for multiple comparisons.

We next assessed whether the greater antitumor effect of F-PRT extended to the AT3 model. In AT3 tumors, F-PRT also produced a greater tumor growth delay than S-PRT and NR (**Fig. 1e**). Tumor-volume endpoint-free survival was improved after F-PRT relative to NR (**Fig. 1f**), and prolonged responses followed by late relapse occurred in 2 of 10 F-PRT-treated mice and in none of the 9 S-PRT-treated mice (**Supplementary Fig. 1b**). Among tumors with measurable exponential growth, F-PRT prolonged tumor doubling time compared with NR, whereas S-PRT did not (**Fig. 1g**).

To determine whether these differences in tumor control were accompanied by changes in the immune microenvironment, tumors were analyzed by multiparametric flow cytometry 7 days after irradiation. In EO771 tumors, F-PRT increased the proportion of CD8+ T cells among CD3+ T cells compared with S-PRT (**Fig. 1h**). In AT3 tumors, F-PRT reduced the proportion of FOXP3+CD25+ regulatory T cells among CD4+ T cells compared with S-PRT (**Fig. 1h**). No additional differences between irradiation modalities were detected among the tumor immune populations analyzed (**Supplementary Fig. 1c**). Splenic immune profiles were also similar between irradiation groups, except for a higher proportion of CD8+IFN-γ+ T cells in AT3-bearing mice after F-PRT than after S-PRT (**Supplementary Fig. 1d**).

Together, these findings show that proton FLASH produces greater tumor growth delay than standard-dose-rate proton irradiation in two heterotopic TNBC models. Complete or prolonged responses were observed only after F-PRT and were accompanied by model-dependent changes in intratumoral T-cell populations: an increase in CD8+ T cells in EO771 and a reduction in regulatory T cells in AT3.

### The enhanced tumor-control effect of proton FLASH is maintained in orthotopic EO771 tumors

Having observed greater tumor control after F-PRT in heterotopic tumors, we next evaluated whether this effect was maintained in orthotopic mammary fat pad models, which more closely recapitulate the tissue-specific tumor microenvironment of breast cancer. EO771 and AT3 tumors received a single 13.5-Gy fraction approximately 10 days after implantation, when they reached a volume of approximately 80 mm³, and were monitored three times per week until endpoint (**Fig. 2a**).

**Fig. 2.**
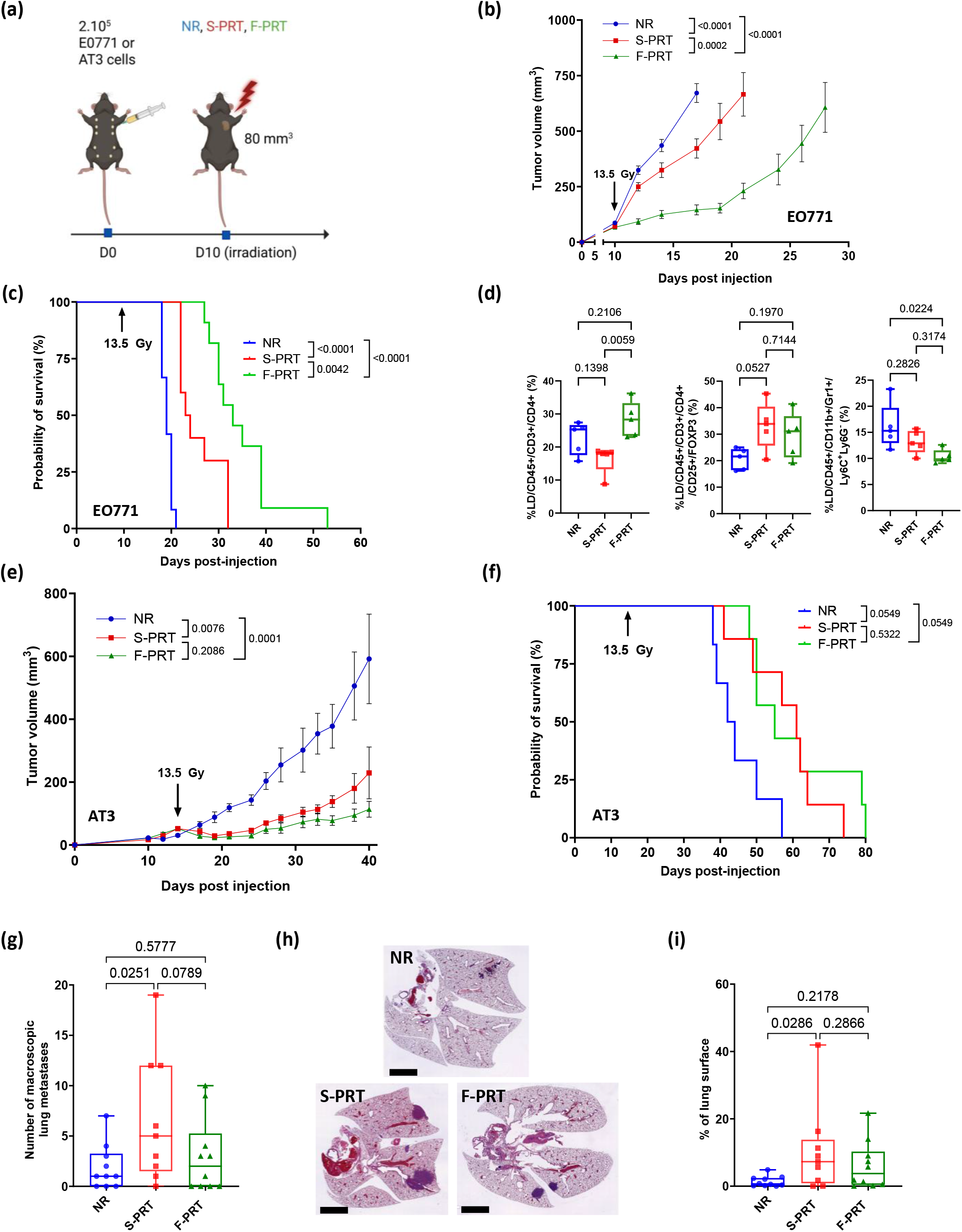
FLASH proton radiotherapy improves tumor control in orthotopic EO771 and AT3 breast tumor models. (a) Experimental design. EO771 or AT3 tumor cells were orthotopically implanted into the second left mammary fat pad. Tumors were irradiated when they reached approximately 80 mm³, 10 days after implantation. Mice were assigned to non-irradiated control (NR), standard dose-rate proton radiotherapy (S-PRT), or FLASH proton radiotherapy (F-PRT). Tumor growth was monitored three times per week until endpoint. (b) Mean tumor growth curves of EO771 tumors after NR, S-PRT, or F-PRT. Group sizes were NR, *n* = 12; S-PRT, *n*= 10; and F-PRT, *n* = 11. Data are shown as mean ± SEM. Tumor growth curves were compared using a mixed-effects model. (c) Kaplan-Meier analysis of tumor-volume endpoint-free survival in the EO771 model, defined as the time to reach 1,000 mm³. Group sizes were NR, *n* = 12; S-PRT, *n* = 10; and F-PRT, *n* = 11. Survival curves were compared using log-rank tests with Holm-Šídák correction for multiple comparisons. (d) Flow cytometry quantification of tumor-infiltrating CD4⁺ T cells, regulatory T cells, and monocytic myeloid-derived suppressor cells in EO771 tumors 7 days after irradiation. Group sizes were *n* = 5 per condition. Boxes show the median and interquartile range, whiskers indicate minimum and maximum values, and each point represents one mouse. Comparisons were performed using ordinary one-way ANOVA followed by Tukey’s multiple-comparison test. (e) Mean tumor growth curves of AT3 tumors after NR, S-PRT, or F-PRT. Group sizes were NR, *n* = 6; S-PRT, *n* = 7; and F-PRT, *n* = 7. Data are shown as mean ± SEM. Tumor growth curves were compared using a mixed-effects model. (f) Kaplan-Meier analysis of tumor-volume endpoint-free survival in the AT3 model, defined as the time to reach 1,000 mm³. Group sizes were NR, *n* = 6; S-PRT, *n* = 7; and F-PRT, *n* = 7. Survival curves were compared using log-rank tests with Holm-Šídák correction for multiple comparisons. (g) Number of macroscopic lung metastases assessed at terminal endpoint in EO771 tumor-bearing mice. Group sizes were NR, *n* = 10; S-PRT, *n* = 9; and F-PRT, *n* = 10. Boxes show the median and interquartile range, whiskers indicate minimum and maximum values, and each point represents one mouse. Comparisons were performed using ordinary one-way ANOVA. (h) Representative hematoxylin and eosin-stained lung sections showing metastatic lesions at terminal endpoint in EO771 tumor-bearing mice. Scale bar, 3 mm. (i) Quantification of metastatic lung involvement on hematoxylin and eosin-stained lung sections, in EO771 tumor-bearing mice, expressed as the proportion of lung surface area occupied by metastases. Group sizes were NR, *n* = 10; S-PRT, *n* = 9; and F-PRT, *n* = 10. Boxes show the median and interquartile range, whiskers indicate minimum and maximum values, and each point represents one mouse. Comparisons were performed using ordinary one-way ANOVA.

In EO771 tumors, both S-PRT and F-PRT delayed tumor growth compared with NR, with a greater effect after F-PRT than after S-PRT (**Fig. 2b** and **Supplementary Fig. 2a**). F-PRT also improved tumor-volume endpoint-free survival compared with S-PRT while both irradiation modalities improved endpoint-free survival relative to NR (**Fig. 2c**). Tumor doubling time was longer after both F-PRT and S-PRT than after NR (**Supplementary Fig. 2b**). We next examined the immune composition of orthotopic EO771 tumors 7 days after irradiation. Immune changes were more limited than those observed in the heterotopic experiments. F-PRT increased the proportion of CD4+ T cells among CD3+ T cells compared with S-PRT, whereas the proportion of CD8+ T cells was comparable between the two irradiation modalities (**Fig. 2d** and **Supplementary Fig. 2c**). The proportion of regulatory T cells among CD4+ T cells was increased after S-PRT compared with NR but not after F-PRT. F-PRT also reduced monocytic myeloid-derived suppressor cells compared with NR (**Fig. 2d**). No additional differences between S-PRT and F-PRT were detected in tumor or splenic immune populations (**Supplementary Fig. 2c,d**). In AT3 tumors, both irradiation modalities also delayed tumor growth compared with NR, with comparable effects for F-PRT and S-PRT (**Fig. 2e** and **Supplementary Fig. 2e)**. Tumor-volume endpoint-free survival and tumor doubling time followed the same pattern, although prolonged responses were observed in 2 of 7 F-PRT-treated mice and 1 of 7 S-PRT-treated mice (**Fig. 2f** and **Supplementary Fig. 2e,f**). The differential effect of dose rate was therefore more evident in EO771 than in AT3 tumors, mirroring the heterotopic experiments. Body-weight trajectories were comparable across the NR, S-PRT, and F-PRT groups in both tumor models, with largely overlapping curves throughout follow-up (**Supplementary Fig. 2g**).

Spontaneous pulmonary metastatic burden was assessed in EO771-bearing mice at the tumor-volume endpoint. Both macroscopic lung nodule counts and the proportion of lung parenchyma occupied by metastases were significantly increased after S-PRT compared with non-irradiated controls, whereas metastatic burden after F-PRT remained comparable to that in non-irradiated controls (**Fig. 2g-i**). F-PRT-treated mice reached the tumor-volume endpoint later than S-PRT-treated mice and therefore had a longer interval during which metastatic dissemination could occur, indicating that the lower metastatic burden was not attributable to shorter follow-up. Together, these findings support a greater antitumor effect of F-PRT in the orthotopic EO771 model, accompanied by selective changes in the tumor immune compartment and a lower spontaneous pulmonary metastatic burden. The consistency of the differential response observed in EO771 across experimental settings provided the basis for subsequent mechanistic studies.

### CD8+ T cells mediate the enhanced tumor control induced by proton FLASH radiotherapy

Because EO771 tumors showed the most consistent difference in tumor control between F-PRT and S-PRT, we next examined the contribution of CD8+ T cells to this response. To complement the relative immune-cell frequencies obtained by flow cytometry, we quantified intratumoral CD8⁺ cell density by immunofluorescence in EO771 tumors collected 2 and 7 days after irradiation. CD8⁺ cell density in EO771 tumors was comparable across all groups at day 2 (**Fig. 3a** and **Supplementary Fig. 3a**). By day 7, however, both irradiated groups showed increased CD8⁺ density relative to non-irradiated controls, with F-PRT producing a significantly greater increase than S-PRT (**Fig. 3b,c**). We next tested whether CD8+ T cells were required for the differential tumor-control effect of F-PRT. EO771-bearing mice received an anti-CD8 depleting antibody or rat IgG isotype control beginning 1 day before irradiation and every 4 days thereafter (**Fig. 3d**). Tumors received a single 13.5-Gy fraction of S-PRT or F-PRT or were left non-irradiated (**Fig. 3d**). Individual tumor growth curves for all six groups are shown in **Supplementary Fig. 3b**, and CD8+ T-cell depletion was confirmed by flow cytometry of peripheral blood (**Supplementary Fig. 3c,d**). In line with the preceding experiments, F-PRT produced greater tumor growth delay and improved tumor-volume endpoint-free survival compared with S-PRT in IgG-treated mice (**Fig. 3e,f**). These differences were no longer observed after CD8+ T-cell depletion (**Fig. 3e,f**). Tumor doubling time was comparable between S-PRT and F-PRT in both IgG-treated and CD8-depleted mice (**Supplementary Fig. 3e**). Together, these findings show that proton FLASH increases intratumoral CD8+ T-cell infiltration compared with standard-dose-rate irradiation and that CD8+ T cells are required for its differential tumor-control effect.

**Fig. 3.**
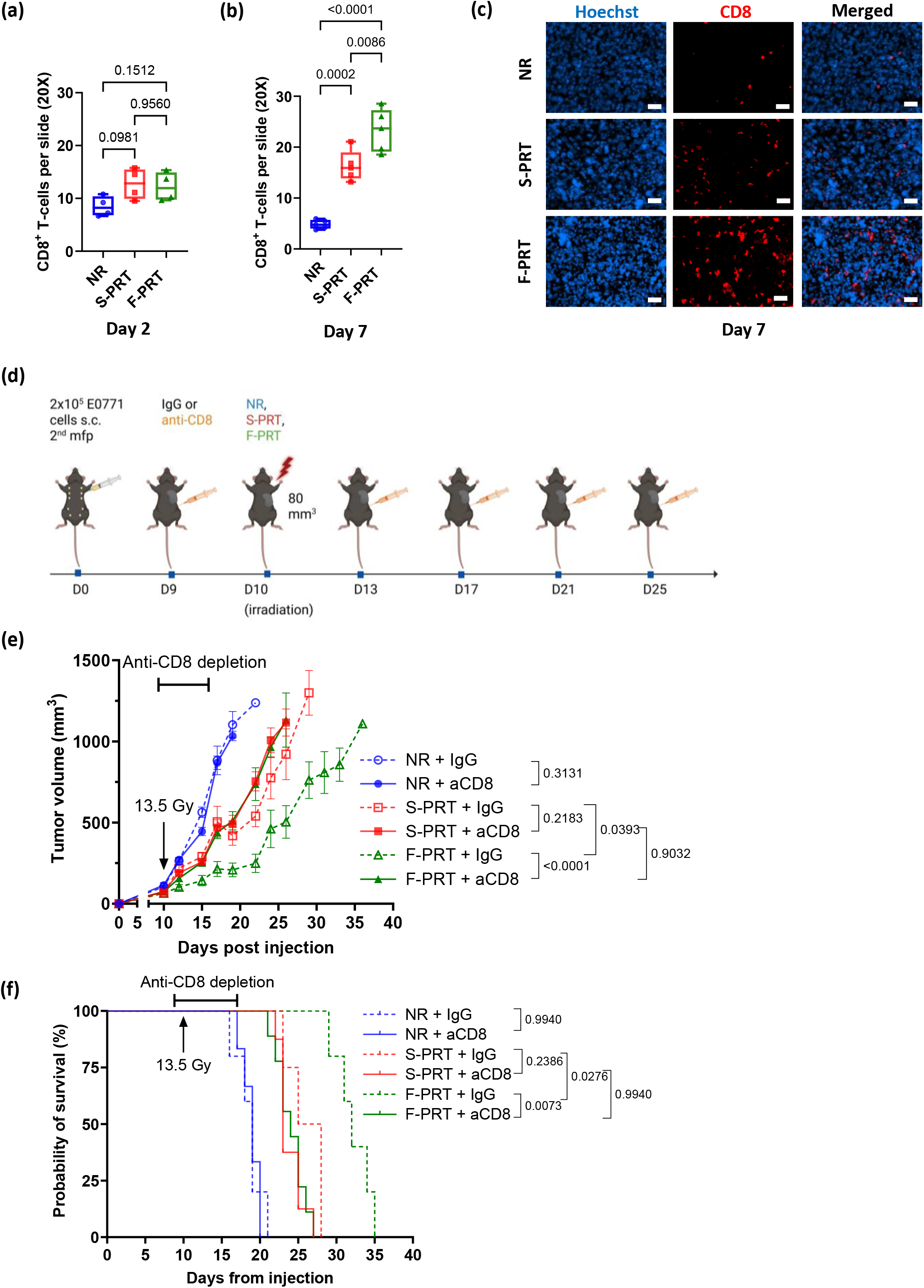
CD8⁺ T-cell depletion abrogates the superior antitumor efficacy of FLASH proton radiotherapy. (a, b) Quantification of CD8⁺ cells in non-irradiated tumors (NR, blue), standard dose-rate proton radiotherapy-treated tumors (S-PRT, red), and FLASH proton radiotherapy-treated tumors (F-PRT, green) at days 2 and 7 post-irradiation, assessed by immunofluorescence and expressed as the number of CD8⁺ cells per 20× field. Tumor samples were obtained from n = 4 mice per group at day 2 and n = 5 mice per group at day 7. For each mouse, 10–15 20× microscopy fields, spatially distributed across the tumor section to provide representative tumor coverage, were quantified and averaged to obtain a single mouse-level value. Box-and-whisker plots show the median, Q1–Q3, and min–max whiskers, and each point represents one mouse. Statistical comparisons were performed using a Kruskal–Wallis test followed by Dunn’s multiple-comparison correction; exact *p* values are shown in the panel. (c) Representative immunofluorescence images of CD8 staining in tumor sections from non-CD8-depleted NR, S-PRT, and F-PRT groups at day 7 post-irradiation. Scale bars, 50 µm. (d) Experimental design of CD8⁺ T-cell depletion in EO771 tumor-bearing mice. Mice received an anti-CD8 depleting antibody or rat IgG isotype control starting one day before irradiation and every 4 days thereafter until endpoint. Tumors were treated with a single 13.5-Gy fraction of S-PRT or F-PRT, or left non-irradiated. Tumor growth and survival were monitored until tumors reached the predefined endpoint volume of 1,000 mm³. (e) Mean tumor growth curves in CD8-depleted mice and rat IgG isotype control mice in the NR, S-PRT, or F-PRT groups. NR is shown in blue, S-PRT in red, and F-PRT in green. Group sizes were NR IgG, *n* = 5; NR anti-CD8, *n* = 6; S-PRT IgG, *n* = 4; S-PRT anti-CD8, *n* = 8; F-PRT IgG, *n* = 5; and F-PRT anti-CD8, *n* = 8. Error bars indicate the SEM. Tumor growth curves were compared using a mixed-effects model. (f) Kaplan-Meier analysis of tumor-volume endpoint-free survival in CD8-depleted mice and rat IgG isotype control mice, defined as the time to reach 1,000 mm³. NR is shown in blue, S-PRT in red, and F-PRT in green. Group sizes were NR IgG, *n* = 5; NR anti-CD8, *n* = 6; S-PRT IgG, *n* = 4; S-PRT anti-CD8, *n* = 8; F-PRT IgG, *n* = 5; and F-PRT anti-CD8, *n* = 8. Survival curves were compared using log-rank tests with Holm–Šídák correction for multiple comparisons; exact *p* values are shown in the panel.

### Proton FLASH radiotherapy promotes early STING–IRF3 signaling and STING-dependent tumor control

Because cGAS-STING-dependent type I interferon signaling promotes radiation-induced dendritic-cell activation and CD8+ T-cell recruitment [59,60], we next examined whether this pathway was differentially activated by F-PRT. STING immunofluorescence showed an increase in STING-positive cells 2 days after F-PRT compared with both NR and S-PRT tumors (**Fig. 4a,b**). By day 7, STING expression was increased in both irradiated groups compared with NR and was comparable between F-PRT and S-PRT, indicating earlier STING induction after F-PRT (**Fig. 4c** and **Supplementary Fig. 4a**).

**Fig. 4.**
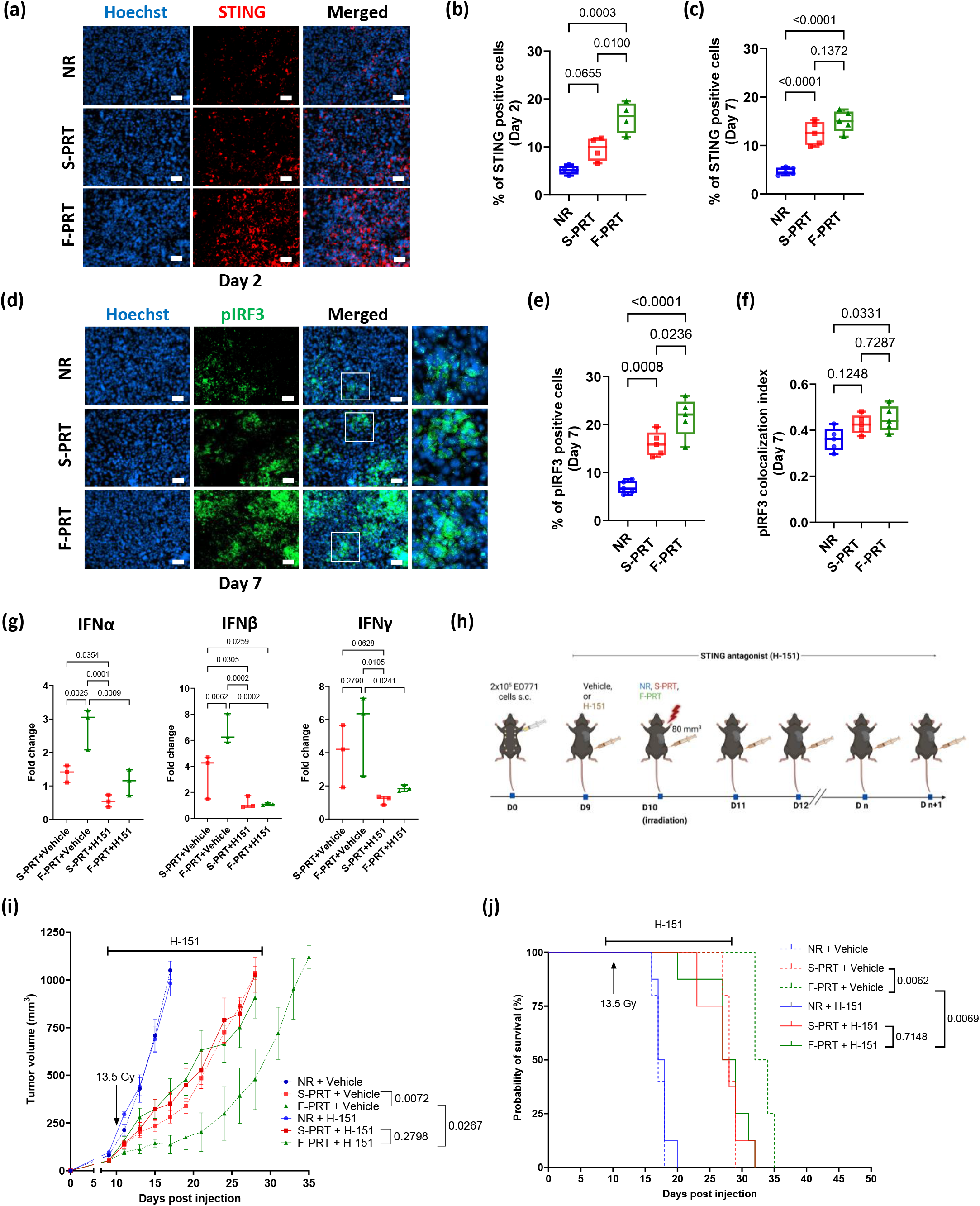
FLASH proton irradiation promotes early STING–IRF3 pathway activation, while STING antagonism abrogates its tumor-control benefit. (a) Representative immunofluorescence staining of STING in EO771 tumors 2 days after irradiation (20× microscopy field). Tumors were analyzed in the non-irradiated control (NR), standard dose-rate proton irradiation (S-PRT), and FLASH proton irradiation (F-PRT) groups. Nuclei were counterstained with Hoechst. The scale bar corresponds to 50 µm. (b, c) Quantification of STING-positive cells in EO771 tumors at days 2 (b) and 7 (c) after irradiation in the NR, S-PRT, and F-PRT groups. For day 2, tumor samples were obtained from n = 4 mice per group, and for day 7 from n = 5 mice per group. For each mouse, 10-15 20× microscopy fields, spatially distributed across the tumor section to provide representative tumor coverage, were quantified and averaged to obtain a single mouse-level value. Boxes show the median and interquartile range, whiskers indicate minimum and maximum values, and each point represents one mouse. Comparisons were performed using ordinary one-way ANOVA with Tukey’s correction for multiple comparisons. (d) Representative immunofluorescence staining of phospho-IRF3 (pIRF3) in EO771 tumors 7 days after irradiation in the NR, S-PRT, and F-PRT groups (20× microscopy field). Nuclei were counterstained with Hoechst. White boxed regions are shown magnified (×3.5) in the right-hand column. The scale bar corresponds to 50 µm. (e, f) Quantification of pIRF3 immunofluorescence in EO771 tumors 7 days after irradiation. Analyses included the proportion of pIRF3-positive cells, irrespective of subcellular localization (e), and the proportion of nuclear pIRF3 signal, defined as pIRF3 colocalized with Hoechst-positive nuclei (f). Tumor samples were obtained from n = 5 mice per group. For each mouse, 10-15 20× microscopy fields, spatially distributed across the tumor section to provide representative tumor coverage, were quantified and averaged to obtain a single mouse-level value. Boxes show the median and interquartile range, whiskers indicate minimum and maximum values, and each point represents one mouse. Comparisons were performed using ordinary one-way ANOVA with Tukey’s correction for multiple comparisons. (c) Multiplex bead-based quantification of IFN-α, IFN-β, and IFN-γ in EO771 tumors 7 days after S-PRT or F-PRT, with or without H-151 treatment. Values are expressed as fold change relative to the non-irradiated (NR) group, set to 1 and not plotted. IFN-α and IFN-β were used as type I interferon markers of STING pathway activation, while IFN-γ was assessed as an immune effector cytokine. Each point represents one mouse (n = 3 per group). Comparisons were performed using ordinary one-way ANOVA. (d) Experimental design of STING antagonism. C57BL/6 mice were orthotopically implanted with 2 × 10⁵ EO771 cells into the left second mammary fat pad. Nine days after tumor implantation, one day before irradiation, mice began daily treatment with the STING antagonist H-151 or vehicle control. When tumors reached approximately 80 mm³, 10 days after implantation, mice were assigned to NR, S-PRT, or F-PRT. H-151 or vehicle administration was continued daily until the predefined tumor endpoint. (e) Mean tumor growth kinetics of EO771 tumors in the NR, S-PRT, and F-PRT groups, with concurrent administration of H-151 or vehicle control. Data are shown as mean ± SEM. NR, S-PRT, and F-PRT are shown in blue, red, and green, respectively. Group sizes were NR + vehicle, n = 5; S-PRT + vehicle, n = 5; F-PRT + vehicle, n = 4; and n = 8 per group for NR + H-151, S-PRT + H-151, and F-PRT + H-151. Tumor growth curves were compared using a longitudinal mixed-effects model. (f) Kaplan-Meier analysis of tumor endpoint-free survival in the NR, S-PRT, and F-PRT groups, with concurrent administration of H-151 or vehicle control. Mice were followed until tumors exceeded the predefined endpoint volume of 1,000 mm³, at which point they were euthanized. NR, S-PRT, and F-PRT are shown in blue, red, and green, respectively. Group sizes were NR + vehicle, n = 5; S-PRT + vehicle, n = 5; F-PRT + vehicle, n = 4; and n = 8 per group for NR + H-151, S-PRT + H-151, and F-PRT + H-151. Survival curves were compared using log-rank tests with Holm-Šídák correction for multiple comparisons.

We next assessed downstream IRF3 activation. At day 2, nuclear pIRF3 colocalization was increased after F-PRT, consistent with early nuclear translocation of activated IRF3, whereas total phosphorylated IRF3 (pIRF3) positivity was comparable across groups (**Supplementary Fig. 4b-d**). By day 7, pIRF3 positivity was increased in both irradiated groups compared with NR and was higher after F-PRT than after S-PRT (**Fig. 4d,e**). Nuclear pIRF3 signal was also increased after F-PRT compared with NR (**Fig. 4f**).

Consistent with enhanced STING–IRF3 activation, tumor concentrations of IFN-α and IFN-β were higher after F-PRT than after S-PRT, and IFN-γ followed the same direction (**Fig. 4g**). The T-cell-recruiting chemokines CXCL9/MIG and CCL5/RANTES showed the same pattern, with the highest levels observed in F-PRT tumors (**Supplementary Fig. 4e**).

We then tested whether STING signaling contributed to the differential antitumor effect of F-PRT. EO771-bearing mice received the covalent STING antagonist H-151 [61] or vehicle beginning 1 day before irradiation and daily thereafter (**Fig. 4h**). Consistent with our earlier findings, F-PRT produced greater tumor growth delay and improved tumor-volume endpoint-free survival relative to S-PRT in vehicle-treated mice. Under STING inhibition (H-151), this advantage was abolished, indicating that the superior efficacy of F-PRT depends on intact STING signaling (**Fig. 4i,j** and **Supplementary Fig. 4f**). Tumor doubling time was comparable between S-PRT and F-PRT in both vehicle- and H-151-treated mice (**Supplementary Fig. 4g,h**).

Under H-151 treatment, the difference in intratumoral CD8+ T-cell infiltration between F-PRT and S-PRT was no longer observed, and interferon and chemokine induction was attenuated across irradiation groups (**Fig. 4g and Supplementary Fig. 4e,i,j**). Differences in pIRF3 positivity and nuclear localization were similarly no longer detected under STING inhibition (**Supplementary Fig. 4k-m**).

Together, these findings show that proton FLASH induces earlier STING-IRF3 activation and greater type I interferon production than standard-dose-rate irradiation. The loss of the differential CD8+ T-cell infiltration and tumor-control effects under pharmacological STING inhibition supports a functional contribution of STING signaling to the proton FLASH response.

### Proton FLASH accelerates tumor response to combined anti-PD-1 and anti-CD40 immunotherapy

Given the increased CD8+ T-cell infiltration after F-PRT and the requirement for CD8+ T cells in its differential antitumor effect, we next examined whether proton FLASH could enhance the response to immunotherapy. Orthotopic EO771 tumors were treated with S-PRT or F-PRT in combination with anti-PD-1 and agonistic anti-CD40, while a non-irradiated group received immunotherapy alone and IgG-treated groups served as controls (**Fig. 5a**).

**Fig. 5.**
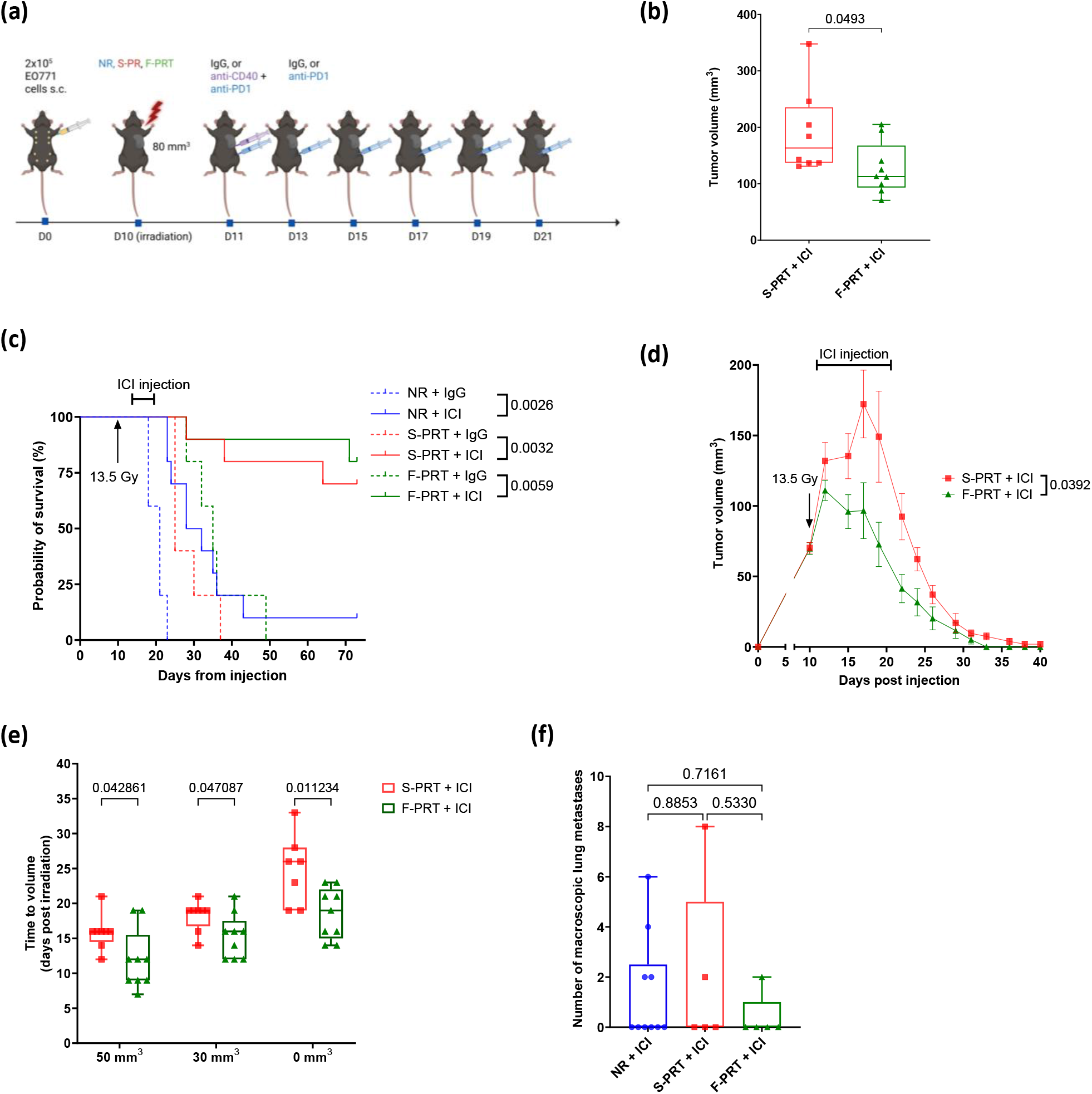
FLASH proton irradiation combined with anti-PD-1 and agonistic anti-CD40 immunotherapy is associated with faster tumor regression than standard dose-rate proton irradiation in an orthotopic EO771 breast cancer model. (a) Experimental design. C57BL/6 mice were orthotopically implanted with 2 × 10⁵ EO771 cells into the left second mammary fat pad. When tumors reached approximately 80 mm³, around 10 days after implantation, mice were either left non-irradiated (NR) or treated with a single 13.5-Gy fraction of standard dose-rate proton irradiation (S-PRT) or FLASH proton irradiation (F-PRT). One day after irradiation, mice received combined immunotherapy consisting of anti-CD40 and anti-PD-1 antibodies for the first injection, followed by five additional injections of anti-PD-1 antibody alone, for a total of six immunotherapy administrations. Tumor growth was monitored longitudinally. (b) Maximum tumor volume reached before regression among responders in the S-PRT plus immunotherapy (S-PRT + ICI; n = 8) and F-PRT plus immunotherapy (F-PRT + ICI; n = 9) groups. S-PRT and F-PRT are shown in red and green, respectively. Boxes show the median and interquartile range, whiskers indicate minimum and maximum values, and each point represents one mouse. Comparisons were performed using Student’s t-test. (c) Kaplan-Meier analysis of tumor endpoint-free survival. Mice were followed until tumors exceeded the predefined endpoint volume of 1,000 mm³, at which point they were euthanized. NR, S-PRT, and F-PRT are shown in blue, red, and green, respectively. Group sizes were n = 5 per group for NR + IgG, S-PRT + IgG, and F-PRT + IgG, and n = 10 per group for NR plus immunotherapy (NR + ICI), S-PRT + ICI, and F-PRT + ICI. Survival curves were compared using log-rank tests with Holm-Sidak correction for multiple comparisons. (d) Tumor growth kinetics among responders after S-PRT (n = 8) or F-PRT (n = 9) combined with immunotherapy, including mice with subsequent late relapse. S-PRT and F-PRT are shown in red and green, respectively. Longitudinal tumor growth curves were compared using a mixed-effects model. (e) Time from irradiation to tumor regression below predefined volume thresholds of 50 mm³, 30 mm³, and 0 mm³ among complete responders in the S-PRT and F-PRT groups. For the 50 mm³ and 30 mm³ thresholds, sample sizes were n = 8 for S-PRT and n = 9 for F-PRT. For the 0 mm³ threshold, sample sizes were n = 7 for S-PRT and n = 9 for F-PRT, as one S-PRT responder did not reach complete tumor regression. S-PRT and F-PRT are shown in red and green, respectively. Boxes show the median and interquartile range, whiskers indicate minimum and maximum values, and each point represents one mouse. Comparisons between S-PRT and F-PRT were performed using Student’s t-tests for each volume threshold. (f) Quantification of pulmonary metastatic burden after treatment. Lung metastases were assessed separately in the right and left lungs, using the number of macroscopic metastatic nodules. Boxes show the median and interquartile range, whiskers indicate minimum and maximum values, and each point represents one mouse. For macroscopic nodule counts, group sizes were NR plus immunotherapy, n = 10; S-PRT plus immunotherapy, n = 5; and F-PRT plus immunotherapy, n = 5, and groups were compared using ordinary one-way ANOVA.

The addition of proton irradiation to immunotherapy produced high rates of tumor regression. Initial regression occurred in 9 of 10 mice after F-PRT and in 8 of 10 mice after S-PRT, compared with 1 of 10 mice receiving immunotherapy alone. Among responders, the maximum tumor volume reached before regression was lower after F-PRT than after S-PRT (**Fig. 5b**). One relapse occurred in each irradiated group, with the relapse in the F-PRT group occurring later than that in the S-PRT group (**Supplementary Fig. 5a)**. At the end of follow-up, durable complete responses were observed in 8 of 10 mice after F-PRT and in 7 of 10 mice after S-PRT. Tumor-volume endpoint-free survival was improved after both F-PRT plus immunotherapy and S-PRT plus immunotherapy compared with immunotherapy alone (**Fig. 5c and Supplementary Fig. 5a**).

Although long-term tumor control was similar after S-PRT and F-PRT, the kinetics of response differed between the two irradiation modalities: longitudinal analysis showed faster tumor regression after F-PRT, with a significant group-by-time interaction between F-PRT and S-PRT (**Fig. 5d**). Consistently, the times to regress below 50 mm³ and 30 mm³ and to complete tumor disappearance were shorter after F-PRT than after S-PRT (**Fig. 5e**). At endpoint, macroscopic pulmonary metastases were absent in 4 of 5 mice after F-PRT plus immunotherapy compared with 3 of 5 after S-PRT, with a lower total nodule count in the F-PRT group (**Fig. 5f**).

To determine whether complete tumor rejection was associated with antitumor memory, relapse-free complete responders were rechallenged with EO771 cells in the contralateral mammary fat pad (**Supplementary Fig. 5b**). All rechallenged mice rejected tumor reimplantation, including 7 mice previously treated with S-PRT plus immunotherapy and 8 mice previously treated with F-PRT plus immunotherapy. After rechallenge, F-PRT plus immunotherapy was associated with significantly higher proportions of CD8⁺ central-memory T cells in draining lymph nodes and of CD4⁺ and CD8⁺ central-memory T cells in the spleen compared with non-irradiated controls. Increases of similar direction were observed after S-PRT plus immunotherapy (**Supplementary Fig. 5c,d**). Neither central-memory nor effector-memory compartments differed between mice previously treated with S-PRT and F-PRT (**Supplementary Fig. 5c,d**), indicating that the two modalities generate comparable systemic memory despite their divergent primary tumor responses.

Together, these findings show that proton irradiation enhances the antitumor activity of combined anti-PD-1 and anti-CD40 treatment. Although both irradiation modalities produced similar rates of durable tumor control and antitumor memory, proton FLASH limited tumor progression before response and accelerated tumor regression compared with standard-dose-rate irradiation.

## Discussion

In this study, proton FLASH radiotherapy produced a greater antitumor effect than standard-dose-rate proton irradiation, most consistently in EO771 tumors. This differential response was observed in both heterotopic and orthotopic EO771 tumors, whereas the distinction between dose-rate conditions was less pronounced in AT3. In EO771 tumors, proton FLASH increased intratumoral CD8+ T-cell accumulation and induced earlier STING expression and IRF3 nuclear activation, followed by greater type I interferon production. The differential tumor-control effect was no longer observed after CD8+ T-cell depletion or pharmacological STING inhibition. Together, these findings support a model in which dose rate modulates antitumor immunity at the same physical dose, with STING–IRF3 signaling and CD8+ T cells contributing to the enhanced response to proton FLASH.

Our findings extend previous evidence that FLASH can induce immune responses distinct from those produced by standard-dose-rate irradiation. In murine non-small cell lung cancer, proton FLASH improved tumor control, increased cytotoxic T-cell infiltration and reduced regulatory T cells and immunosuppressive myeloid populations [47]. In medulloblastoma, proton FLASH promoted proinflammatory macrophage states and increased T-cell infiltration [48]. Electron FLASH has also been associated with preservation of interferon signaling and T-cell-recruiting chemokines in TNBC models despite similar tumor control to standard-dose-rate irradiation [62]. These studies consistently implicate interferon signaling, T-cell recruitment and myeloid remodeling in the response to FLASH irradiation, and are complemented by evidence that FLASH improves the efficacy of PD-1 blockade in ovarian cancer models [49] and spares circulating lymphocytes relative to standard dose rates [46,63]. Whether these immune effects translate into greater tumor control likely depends on the irradiation conditions and tumor model; in our study, F-PRT delayed tumor growth in both EO771 and AT3 tumors, but the effect was stronger and accompanied by more pronounced immune changes in EO771. Previous reports of greater sensitivity to PD-1 blockade in EO771 than in AT3 [58] support baseline immune responsiveness as a potential determinant of the differential effects of FLASH. The greater FLASH effect observed in the more checkpoint-responsive EO771 model suggests a potential parallel between determinants of sensitivity to FLASH and to immune checkpoint inhibition, with pre-existing capacity for T-cell-mediated antitumor immunity possibly influencing both [12,64].

The STING–IRF3 response observed here is consistent with established links between radiation-induced DNA damage and innate immune sensing [65]. Cytosolic DNA sensing through cGAS-STING can couple irradiation to type I interferon production, supporting dendritic-cell cross-priming and antitumor CD8+ T-cell responses [59,66]. This response is not necessarily proportional to dose since induction of the DNA exonuclease TREX1 above a cell-dependent threshold can limit cytosolic DNA accumulation and attenuate radiation-induced immunogenicity [66]. Whether FLASH alters this balance has not previously been tested in tumors. To our knowledge, the present study is the first to show that dose-rate-dependent changes in STING–IRF3 signaling within an irradiated tumor. Notably, the effect we observed was kinetic rather than purely quantitative: at an identical physical dose, F-PRT advanced the timing of STING induction and IRF3 nuclear translocation, and by day 7 STING expression had converged between the two dose rates. This pattern is more consistent with dose rate altering how rapidly cytosolic DNA becomes available for sensing than with a change in the total amount of damage generated, and it offers a testable explanation for why a difference in tumor control emerges despite identical delivered dose. Evidence from normal tissues further indicates that FLASH does not uniformly enhance this pathway; in mice receiving abdominal X-ray irradiation with PD-L1 blockade, FLASH reduced cytosolic DNA accumulation and cGAS–STING–IRF3 activation in intestinal crypts, limiting CD8+ T-cell-associated, gasdermin E-mediated pyroptosis while preserving antitumor efficacy [67]. Together with our findings, these observations suggest that the direction and consequences of dose-rate-dependent innate immune signaling may depend on the tissue and treatment context. If this contrast between normal tissue and tumor is confirmed, it would be therapeutically favorable, since attenuated cGAS-STING-driven pyroptosis in irradiated normal tissue alongside enhanced STING–IRF3 signaling in the tumor would widen the therapeutic window through the same pathway that mediates the antitumor effect. Beyond type I interferon signaling, IFN-γ can also contribute to radiation-induced antitumor immunity by enhancing MHC class I expression and inducing chemokines that support T-cell recruitment; genetic loss of IFN-γ also abolished radiation-mediated tumor control in a murine colon carcinoma model [68,69]. These studies provide a mechanistic framework linking innate DNA sensing to adaptive antitumor immunity. Our findings support a role for STING signaling and CD8+ T cells in the enhanced antitumor response to proton FLASH, providing a basis for further investigation of upstream DNA sensing and downstream interferon-mediated effector mechanisms.

The combination of FLASH with immunotherapy may be particularly relevant in high-risk TNBC. Pembrolizumab combined with neoadjuvant chemotherapy and continued after surgery improves event-free and overall survival [18], yet patients with residual invasive disease remain at substantial risk of recurrence [9]. Radiotherapy capable of increasing immune-cell recruitment and reducing local immunosuppression could therefore strengthen perioperative treatment in this population. In our study, anti-PD-1 combined with agonistic anti-CD40 – a pairing that engages both the T-cell and the myeloid compartment and that cooperates with radiotherapy [70–72] – produced high rates of durable complete response with either proton irradiation modality. Compared with standard-dose-rate irradiation, proton FLASH reduced the maximum tumor volume reached before regression and accelerated tumor shrinkage, indicating a faster antitumor response during combined immunotherapy. All relapse-free complete responders rejected tumor rechallenge, consistent with durable antitumor memory after both irradiation modalities. These findings support proton FLASH as an immunologically active partner for immunotherapy and provide a rationale for its evaluation in the preoperative setting, where pathological response and treatment-induced changes in the tumor immune microenvironment could be assessed directly.

The 13.5 Gy single fraction provides a clinically interpretable reference within the progressive reduction in fraction number used in breast radiotherapy. Assuming an α/β ratio of 5 Gy [73], 13.5 Gy in one fraction corresponds to a biologically effective dose of 50.0 Gy₅, compared with 53.0 Gy₅ for the standard FAST-Forward regimen of 26 Gy in five fractions [54]. These estimates provide clinical context rather than direct biological equivalence, as the linear-quadratic model is not validated for high single doses and does not account for dose-rate effects [73–75]. Single-fraction preoperative partial-breast irradiation has already been evaluated in phase I and II studies of selected early-stage breast cancers [76–79], demonstrating the feasibility of irradiating the intact tumor and assessing pathological and biological responses in the surgical specimen. This setting may therefore provide a clinically relevant framework for evaluating FLASH-induced immune activation, including in patients undergoing mastectomy in whom the irradiated tumor could subsequently be analyzed directly. Beyond localized disease, FLASH may also be relevant in the metastatic setting, where the ability to deliver ablative radiation is often constrained by adjacent organs at risk. The durable tumor rejection observed after rechallenge in mice treated with radiotherapy plus immunotherapy further supports investigation of whether FLASH can enhance systemic antitumor immune responses when combined with immunotherapy. Clinical translation will also require proton FLASH delivery techniques that provide adequate target coverage and normal-tissue sparing. Clinical experience with proton FLASH has so far relied on single transmission beams, including in the FAST-01 and FAST-02 trials for painful bone metastases [43,80]. Tangential transmission fields may be feasible for selected whole-breast treatments [81], whereas more complex anatomies or regional nodal irradiation will require conformal delivery techniques that maintain ultra-high dose rates while ensuring adequate target coverage and limiting exit dose. Patient-specific three-dimensional range modulators, including systems such as HEDGEHOG [82], may enable conformal single-energy FLASH delivery in these settings, as may optimized pencil-beam scanning strategies that maintain ultra-high three-dimensional dose rates while conforming dose to the target [83–85]. While the present study provides evidence for a differential biological response to FLASH proton irradiation, several aspects of the experimental design warrant further consideration. The study evaluated two syngeneic tumor models using a single radiation dose of 13.5 Gy delivered in one fraction and a single proton transmission-beam configuration. Further studies across additional tumor models, dose and fractionation schedules, and clinically relevant beam configurations will be needed to establish the generalizability of these findings. Mechanistic experiments were performed mainly in EO771 tumors, and the biological basis for the different responses of EO771 and AT3 was not directly investigated. H-151 was administered systemically, preventing identification of the cellular compartment in which STING signaling contributed to the response; genetic approaches using Sting1-deficient mice or conditional deletion in tumor or myeloid cells would be required to resolve this. Upstream cGAS activation, cytosolic double-stranded DNA and micronucleus formation were not directly measured. Immune analyses were restricted to selected markers and two time points; cytokine, chemokine and pulmonary metastasis findings should therefore be regarded as exploratory and hypothesis-generating. Normal-tissue toxicity was not assessed in these animals; the present study therefore demonstrates improved tumor control rather than a directly measured improvement in therapeutic ratio, although normal-tissue sparing by proton FLASH on this platform, and by photon FLASH in other systems, has been reported previously [38,39,86]. Lungs were collected at the tumor-volume endpoint, which was reached later in F-PRT-treated mice; the lower metastatic burden in this group was therefore observed despite a longer interval at risk, but the analysis was not designed to control for time at risk. Further studies should evaluate fractionated and conformal FLASH delivery and integrate tumor response, immune biomarkers and normal-tissue effects in the same animals.

Proton FLASH produced a greater antitumor effect than standard-dose-rate proton irradiation, with the largest effect in EO771 tumors. Mechanistically, this response required STING signaling and CD8⁺ T cells, and was accompanied by earlier IRF3 activation, increased type I interferon production and greater intratumoral CD8⁺ T-cell infiltration. Dose rate is therefore not merely a determinant of normal-tissue sparing but an independent modulator of tumor immunogenicity. This establishes an immunological rationale for proton FLASH that extends beyond its established therapeutic window advantage and warrants translational evaluation in breast cancer.

## Acknowledgements

The irradiations were performed by the Cell and Animal Radiation Core Facility (RRID:SCR_022377) at the University of Pennsylvania Perelman School of Medicine. We would like to thank the in-house Ion Beam Applications (IBA) physics team for facilitating the preclinical experiments. We thank the members of the Penn Vet Comparative Pathology Core (RRID:SCR_022438) and Histology Laboratory, in particular Esha Banerjee and Lacey Hall Kirby for their assistance with the histology preparations. The veterinary pathologists performing the histopathological analysis are partially supported by the Abramson Cancer Center Support Grant (P30 CA016520). The scanner used for whole slide imaging and the image analysis software was supported by an NIH Shared Instrumentation Grant (S10 OD023465-01A1). The authors also thank the University of Pennsylvania Diabetes Research Center for the use of the Biomarkers Core (P30-DK19525).

## Author contributions

P.L.: Conceptualization, Investigation, Data curation, Formal analysis, Visualization, Writing – original draft, Writing – review & editing. I.P.: Investigation, Data curation, Formal analysis, Writing – review & editing. C.C.: Investigation, Data curation, Writing – review & editing. K.K.: Investigation, Data curation, Writing – review & editing. M.K.: Methodology, Software, Validation, Writing – review & editing. J.S.: Methodology, Investigation, Validation, Writing – review & editing. K.S.: Investigation, Data curation. I.M.B.: Investigation, Data curation. J.M.: Resources, Writing – review & editing. R.A.G.: Writing – review & editing. W.B.: Formal analysis, Methodology, Writing – review & editing. E.B.: Writing – review & editing. C.-A.A.: Investigation, Formal analysis, Writing –review & editing. C.K.: Resources, Funding acquisition, Writing – review & editing. E.D.: Methodology, Investigation, Validation, Writing – review & editing. N.T.: Resources, Writing – review & editing. I.I.V.: Conceptualization, Methodology, Supervision, Project administration, Funding acquisition, Writing – original draft, Writing – review & editing. All authors read and approved the final version of the manuscript.

## Data Availability

Research data are stored in an institutional repository and will be shared upon request to the corresponding author.

