## Supplementary figures and images for "Proton FLASH radiotherapy enhances control of triple-negative breast cancer through STING–IRF3 and CD8+ T-cell immunity"

### Supplementary Figure 1

#### Supplementary Figure 1

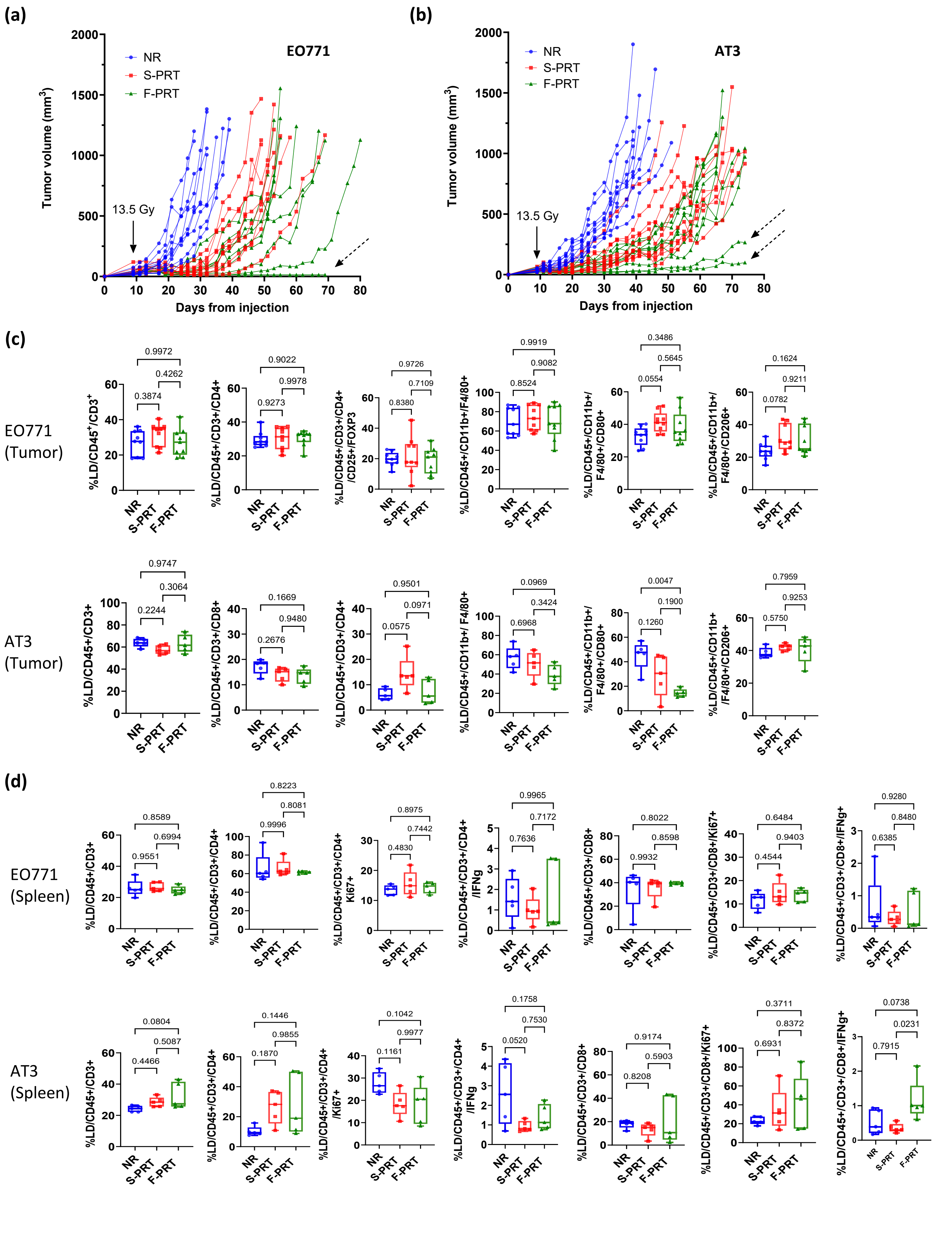

### Supplementary Figure 2

#### Supplementary Figure 2

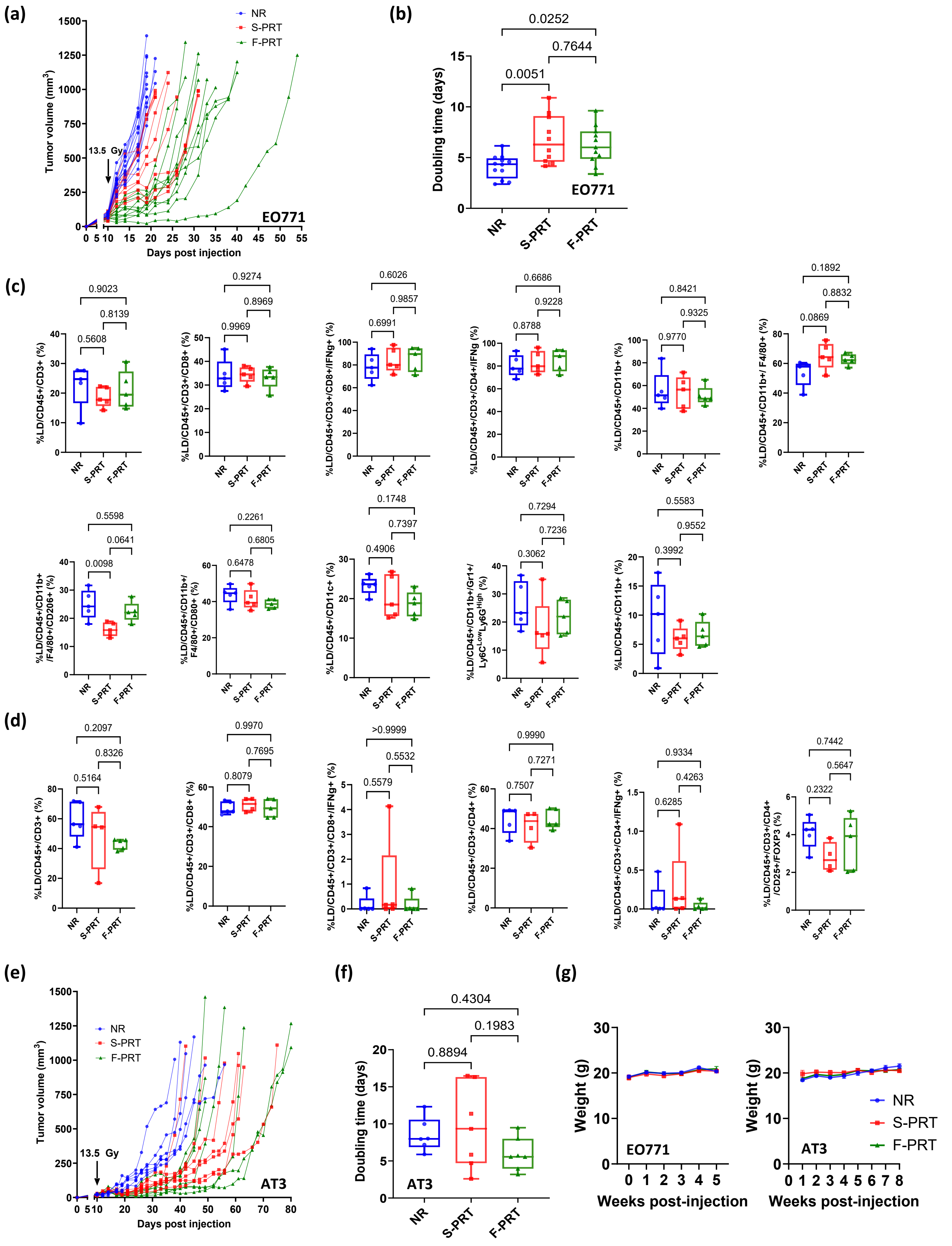

### Supplementary Figure 3

#### Supplementary Figure 3

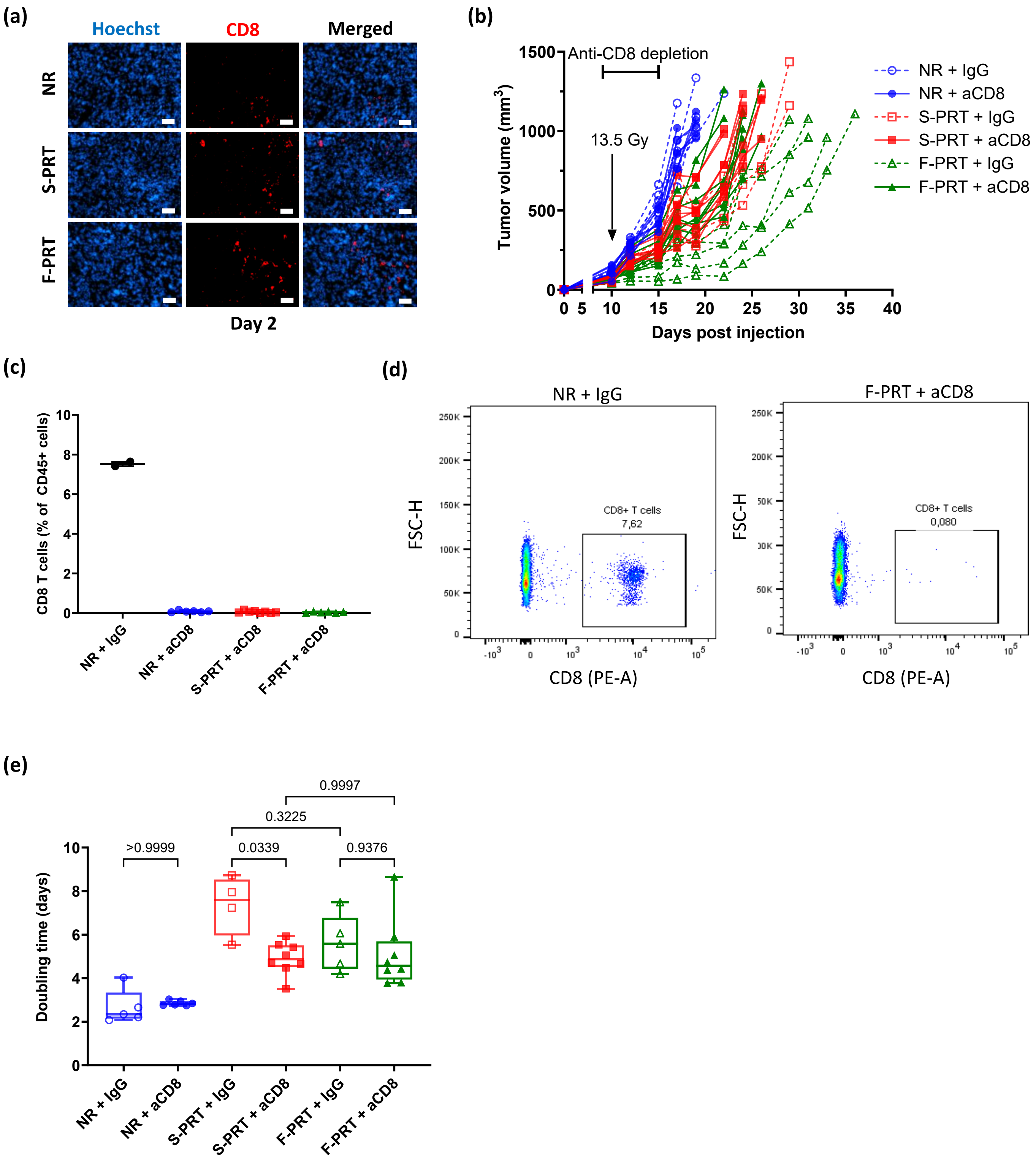

### Supplementary Figure 4

#### Supplementary Figure 4

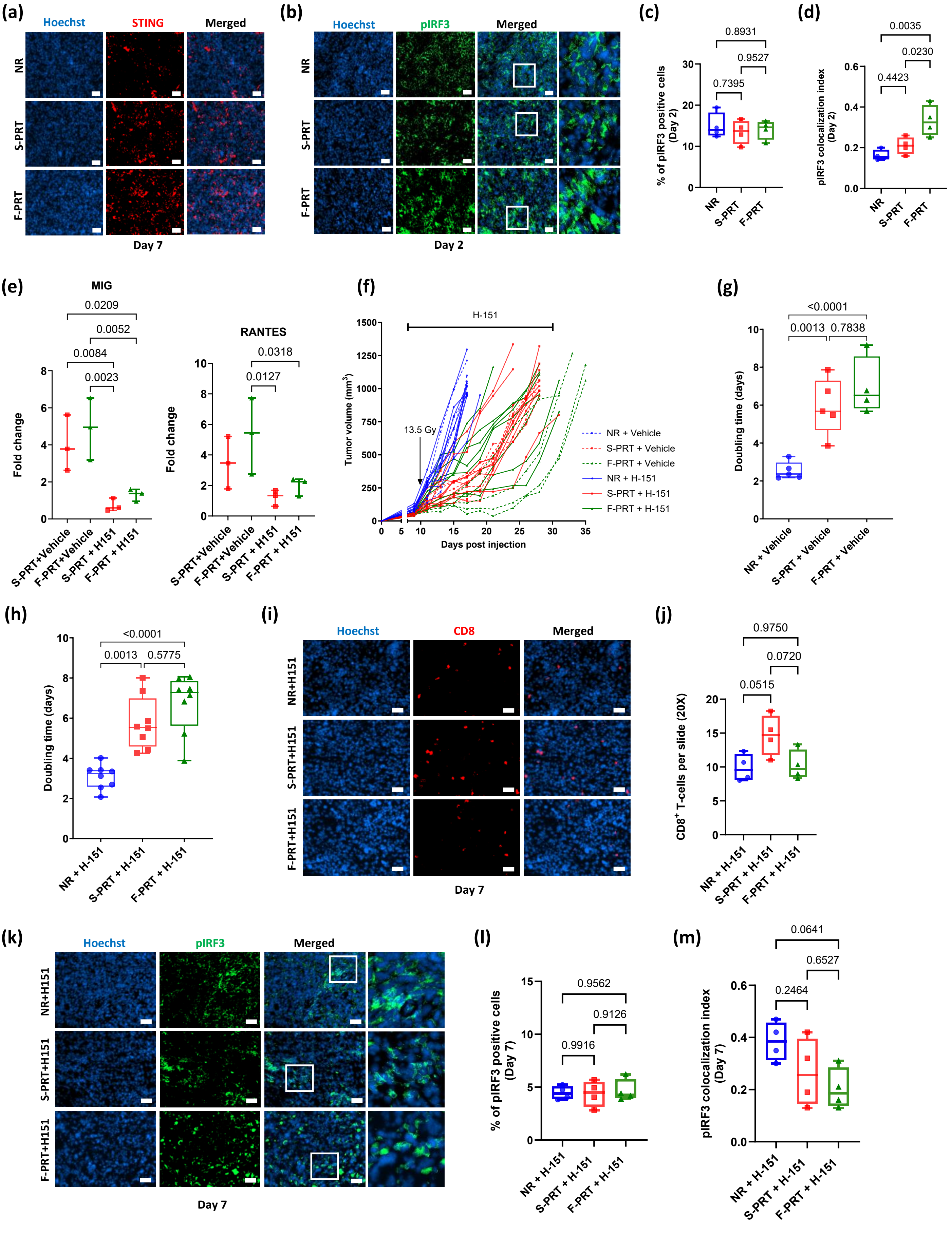

### Supplementary Figure 5

##### Supplementary Figure 5

(a)

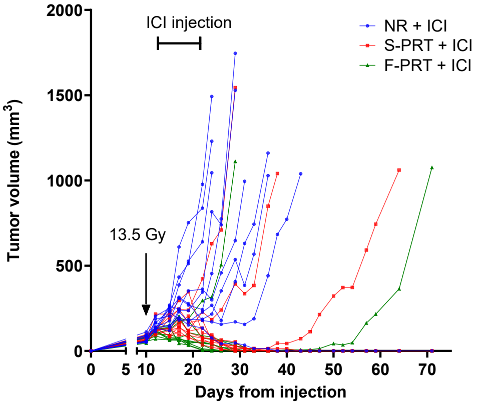


(b)

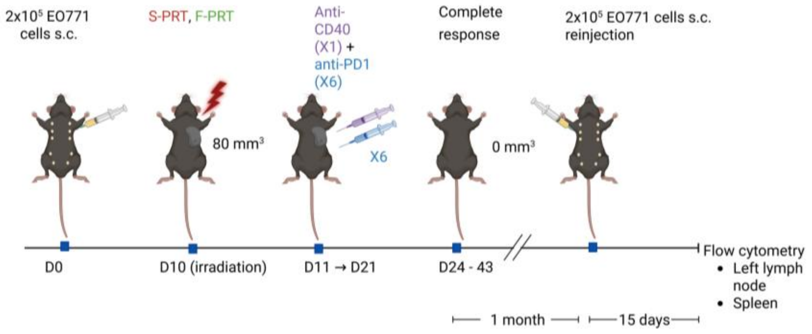


(c)

Lymph node

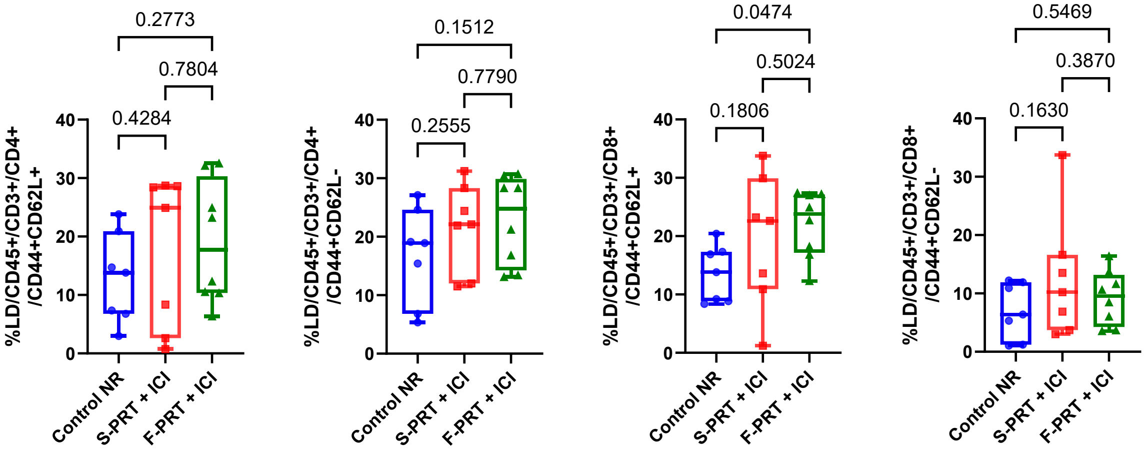


(d)

Spleen

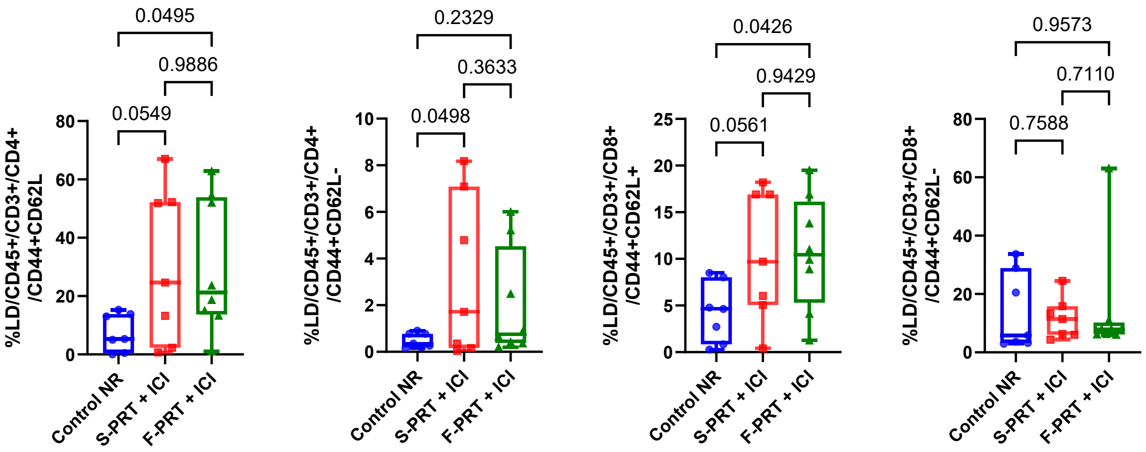

### Supplementary Figure 6

#### Supplementary Figure 6

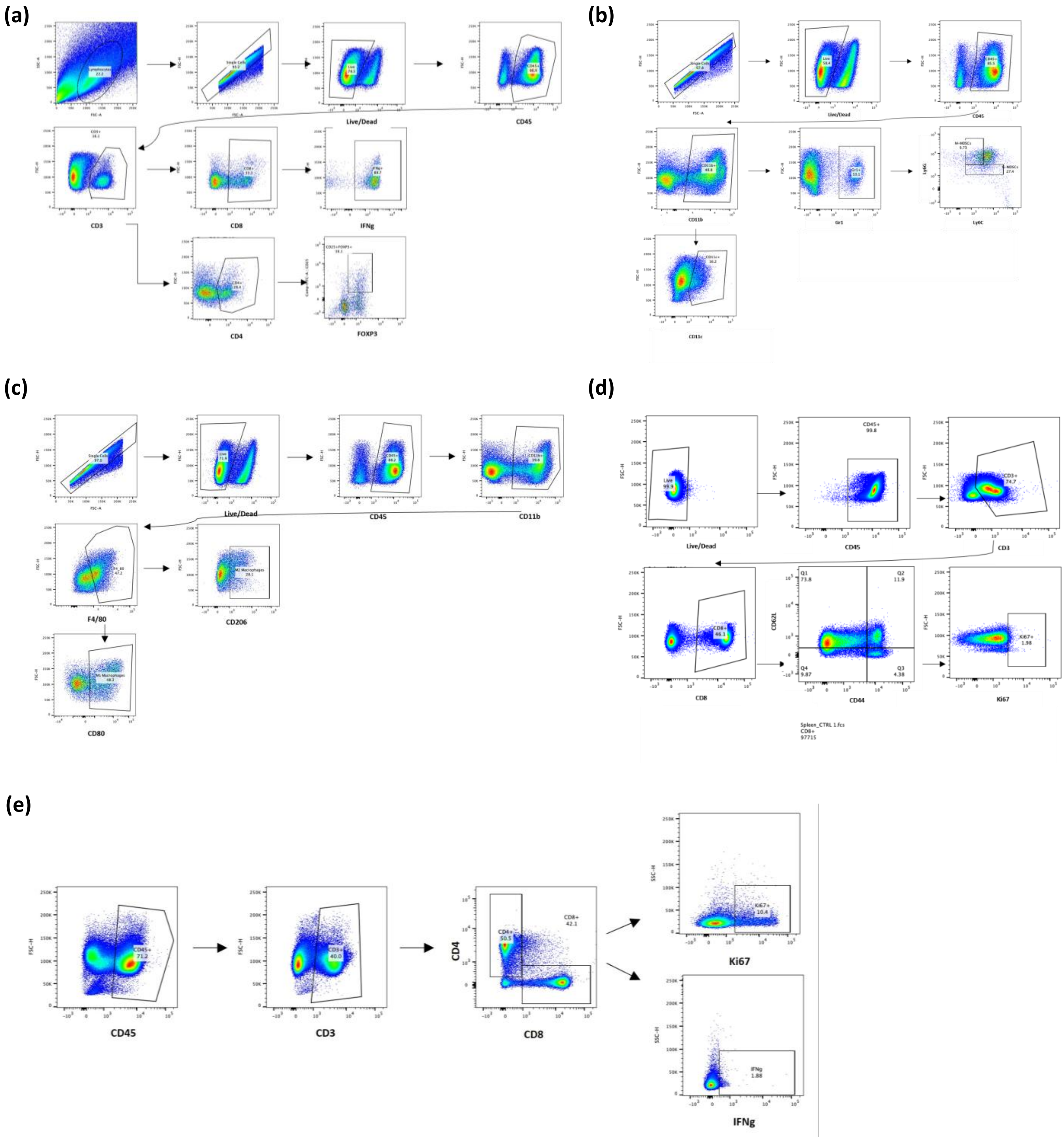
