## Supplementary Figure legends for "Proton FLASH radiotherapy enhances control of triple-negative breast cancer through STING–IRF3 and CD8+ T-cell immunity"

**Supplementary Fig. 1. Individual tumor growth, splenic immune profiling, and additional tumor microenvironment analyses in heterotopic TNBC models.**

**(a,b)** Individual tumor growth curves of EO771 **(a)** and AT3 **(b)** tumors in the non-irradiated control (NR), standard dose-rate proton irradiation (S-PRT), and FLASH proton irradiation (F-PRT) groups. Each curve represents one mouse. Durable tumor responses followed by late relapse are highlighted by dashed arrows. In panel **(a)**, a single dashed arrow indicates two F-PRT-treated EO771 mice with overlapping complete response curves. In panel **(b)**, two dashed arrows indicate two F-PRT-treated AT3 mice with durable tumor responses followed by late relapse.

**(c)** Additional flow cytometry analysis of the tumor microenvironment in EO771 and AT3 tumors 7 days after irradiation, showing immune cell populations not presented in Figure 1. Boxes show the median and interquartile range, whiskers indicate minimum and maximum values, and each point represents one mouse. For EO771 tumors, sample sizes were n = 9 for NR, n = 9 for S-PRT, and n = 9 for F-PRT. For AT3 tumors, sample sizes were n = 5 for NR, n = 5 for S-PRT, and n = 5 for F-PRT. Comparisons were performed using ordinary one-way ANOVA with Tukey's correction for multiple comparisons.

**(d)** Flow cytometry analysis of immune cell populations in the spleen of mice bearing EO771 or AT3 tumors, performed 7 days after irradiation. Boxes show the median and interquartile range, whiskers indicate minimum and maximum values, and each point represents one mouse. Sample sizes were n = 5 for NR, n = 5 for S-PRT, and n = 5 for F-PRT in both tumor models. Comparisons were performed using ordinary one-way ANOVA with Tukey's correction for multiple comparisons.

**Supplementary Fig. 2. Individual tumor growth kinetics, immune profiling, and tumor doubling-time analyses in EO771 and AT3 tumor-bearing mice.**

**(a)** Individual tumor growth curves for each EO771 tumor-bearing mouse after NR, S-PRT, or F-PRT. Tumor volumes were monitored three times per week until tumors reached the predefined endpoint volume of 1,000 $mm^3$. Group sizes were NR, $n = 12$; S-PRT, $n = 10$; and F-PRT, $n = 11$.

**(b)** Tumor doubling time in the EO771 model after NR, S-PRT, or F-PRT. Group sizes were NR, $n = 12$; S-PRT, $n = 10$; and F-PRT, $n = 11$. Doubling time was estimated by fitting an exponential growth model to tumor-volume measurements between 100 and 1,000 $mm^3$. Boxes show the median and interquartile range, whiskers indicate minimum and maximum values, and each point represents one mouse. Comparisons were performed using ordinary one-way ANOVA followed by Tukey's multiple-comparison test.

**(c)** Additional flow cytometry characterization of the EO771 tumor microenvironment 7 days after irradiation. Group sizes were NR, $n = 5$; S-PRT, $n = 5$; and F-PRT, $n = 5$. Boxes show the median and interquartile range, whiskers indicate minimum and maximum values, and each point represents one mouse. Comparisons were performed using ordinary one-way ANOVA followed by Tukey's multiple-comparison test.

**(d)** Flow cytometry characterization of splenic immune populations 7 days after irradiation in EO771 tumor-bearing mice. Group sizes were NR, $n = 5$; S-PRT, $n = 4\text{-}5$; and F-PRT, $n = 5$. Boxes show the median and interquartile range, whiskers indicate minimum and maximum values, and each point represents one mouse. Comparisons were performed using ordinary one-way ANOVA followed by Tukey's multiple-comparison test.

**(e)** Individual tumor growth curves for each AT3 tumor-bearing mouse after NR, S-PRT, or F-PRT. Tumor volumes were monitored three times per week until tumors reached the predefined endpoint volume of 1,000 $mm^3$. Group sizes were NR, $n = 6$; S-PRT, $n = 7$; and F-PRT, $n = 7$.

**(f)** Tumor doubling time in the AT3 model after NR, S-PRT, or F-PRT. Group sizes were NR, $n = 6$; S-PRT, $n = 7$; and F-PRT, $n = 7$. Doubling time was estimated by fitting an exponential growth model to tumor-volume measurements between 100 and 1,000 $mm^3$. Boxes show the median and interquartile range, whiskers indicate minimum and maximum values, and each point represents one mouse. Comparisons were performed using ordinary one-way ANOVA followed by Tukey's multiple-comparison test.

**(g)** Body weight of EO771- and AT3-bearing mice following irradiation. Group sizes for EO771-bearing mice were NR, *n* = 12; S-PRT, *n* = 10; and F-PRT, *n* = 11. Group sizes for AT3-bearing mice were NR, *n* = 6; S-PRT, *n* = 7; and F-PRT, *n* = 7. Data are expressed in grams as mean ± SEM. NR, S-PRT, and F-PRT groups are shown in blue, red, and green, respectively.

**Supplementary Fig. 3. Individual tumor growth kinetics and flow cytometry validation of CD8$^+$ T-cell depletion in EO771 tumor-bearing mice.**

**(a)** Representative immunofluorescence images of CD8 staining in tumor sections from non-CD8-depleted NR, S-PRT, and F-PRT groups at day 2 post-irradiation. Scale bars, 50 µm.

**(b)** Individual tumor growth curves for each EO771 tumor-bearing mouse across the six experimental conditions. Mice received either rat IgG isotype control (IgG) or an anti-CD8 depleting antibody (aCD8) starting one day before irradiation and every 4 days thereafter until endpoint. Tumors were treated with a single 13.5-Gy fraction of standard dose-rate proton radiotherapy (S-PRT) or FLASH proton radiotherapy (F-PRT), or left non-irradiated (NR). Tumor volumes were monitored over time until tumors reached the predefined endpoint volume of 1,000 mm$^3$. Group sizes were NR IgG, *n* = 5; NR anti-CD8, *n* = 6; S-PRT IgG, *n* = 4; S-PRT anti-CD8, *n* = 8; F-PRT IgG, *n* = 5; and F-PRT anti-CD8, *n* = 8.

**(c)** Flow cytometry quality control of CD8$^+$ T-cell depletion in EO771 tumor-bearing mice receiving anti-CD8 depleting antibody. CD8$^+$ T cells were quantified in the three CD8-depleted treatment groups, NR (n=6), S-PRT (n=8), and F-PRT (n=6), as the proportion of CD8$^+$ cells among total CD45$^+$ immune cells.

**(d)** Representative flow cytometry gating of CD8$^+$ cell populations among CD45$^+$ immune cells. Representative plots are shown for an NR IgG control mouse and an F-PRT anti-CD8-depleted mouse, illustrating effective CD8$^+$ T-cell depletion.

**(e)** Tumor doubling time across treatment groups in CD8-depleted mice and rat IgG isotype control mice. For each mouse, doubling time was estimated by fitting an exponential growth model to tumor-volume measurements between 100 and 1,000 $mm^3$ NR is shown in blue, S-PRT in red, and F-PRT in green. Group sizes were NR IgG, $n = 5$; NR anti-CD8, $n = 6$; S-PRT IgG, $n = 4$; S-PRT anti-CD8, $n = 8$; F-PRT IgG, $n = 5$; and F-PRT anti-CD8, $n = 8$. Box-and-whisker plots show the median, Q1–Q3, and min–max whiskers. Statistical comparisons were performed using ordinary one-way ANOVA with correction for multiple comparisons. Exact *p* values are shown in the panel.

**Supplementary Fig. 4. Additional characterization of STING–pIRF3 signaling, chemokine induction, CD8 infiltration, and tumor response after STING antagonism.**

**(a)** Representative immunofluorescence staining of STING in EO771 tumors 7 days after irradiation in the NR, S-PRT, and F-PRT groups. Nuclei were counterstained with Hoechst.

**(b)** Representative immunofluorescence staining of phospho-IRF3 (pIRF3) in EO771 tumors 2 days after irradiation. Tumors were analyzed in the non-irradiated control (NR), standard dose-rate proton radiotherapy (S-PRT), and FLASH proton radiotherapy (F-PRT) groups. Nuclei were counterstained with Hoechst. White boxed regions are shown magnified (×3.5) in the right-hand column. The scale bar corresponds to 50 μm.

**(c, d)** Quantification of pIRF3 immunofluorescence in EO771 tumors 2 days after irradiation, including the proportion of pIRF3-positive cells (**c**) and the proportion of nuclear pIRF3 signal (**d**), defined as pIRF3 colocalized with Hoechst-positive nuclei. Tumor samples were obtained from n = 4 mice per group. For each mouse, 10-15 20× microscopy fields, spatially distributed across the tumor section to provide representative tumor coverage, were quantified and averaged to obtain a single mouse-level value. Boxes show the median and interquartile range, whiskers indicate minimum and maximum values, and each point represents one mouse. Comparisons were performed using ordinary one-way ANOVA with Tukey's correction for multiple comparisons; exact *p* values are shown in the panel.

**(e)** Multiplex bead-based quantification of MIG/CXCL9 and RANTES/CCL5 in EO771 tumors after S-PRT or F-PRT, with concurrent administration of H-151 or vehicle control. Values are expressed as fold change relative to the

NR vehicle-treated group, set to 1. Each point represents one mouse. Comparisons were performed using ordinary one-way ANOVA; exact *p* values are shown in the panel.

**(f)** Individual tumor growth curves of EO771 tumors in the NR, S-PRT, and F-PRT groups, with concurrent administration of the STING antagonist H-151 or vehicle control. Each curve represents one mouse. Group sizes were NR + vehicle, n = 5; S-PRT + vehicle, n = 5; F-PRT + vehicle, n = 4; and n = 8 per group for NR + H-151, S-PRT + H-151, and F-PRT + H-151.

**(g, h)** Tumor doubling time in the NR, S-PRT, and F-PRT groups, with concurrent administration of H-151 (**g**) or vehicle control (**h**). Doubling time was calculated by fitting an exponential growth model to tumor-volume measurements above 100 $mm^3$. Boxes show the median and interquartile range, whiskers indicate minimum and maximum values, and each point represents one mouse. Group sizes were NR + vehicle, n = 5; S-PRT + vehicle, n = 5; F-PRT + vehicle, n = 4; and n = 8 per group for NR + H-151, S-PRT + H-151, and F-PRT + H-151. Comparisons were performed using ordinary one-way ANOVA followed by Tukey's multiple-comparison test; exact *p* values are shown in the panel.

**(i)** Representative immunofluorescence staining of CD8 in EO771 tumors 7 days after irradiation in mice treated with H-151. Tumors were analyzed in the NR, S-PRT, and F-PRT groups. Nuclei were counterstained with Hoechst.

**(j)** Quantification of CD8-positive cells in EO771 tumors 7 days after irradiation in mice treated with H-151. Tumor samples were obtained from n = 4 mice per group. For each mouse, 10-15 20× microscopy fields, spatially distributed across the tumor section to provide representative tumor coverage, were quantified and averaged to obtain a single mouse-level value. Boxes show the median and interquartile range, whiskers indicate minimum and maximum values, and each point represents one mouse. Comparisons were performed using ordinary one-way ANOVA followed by Tukey's multiple-comparison test; exact *p* values are shown in the panel.

**(k)** Representative immunofluorescence staining of phospho-IRF3 (pIRF3) in EO771 tumors 7 days after irradiation in H-151-treated mice from the NR, S-PRT, and F-PRT groups. Nuclei were counterstained with Hoechst. White boxed regions are shown magnified (×3.5) in the right-hand column. The scale bar corresponds to 50 µm.

**(l, m)** Quantification of pIRF3 immunofluorescence in EO771 tumors 7 days after irradiation in mice treated with H-151, including the proportion of pIRF3-positive cells (**l**) and the proportion of nuclear pIRF3 signal (**m**), defined as pIRF3 colocalized with Hoechst-positive nuclei. Tumor samples were obtained from n = 4 mice per group. For each mouse, 10-15 20× microscopy fields, spatially distributed across the tumor section to provide representative tumor coverage, were quantified and averaged to obtain a single mouse-level value. Boxes show the median and interquartile range, whiskers indicate minimum and maximum values, and each point represents one mouse. Comparisons were performed using ordinary one-way ANOVA followed by Tukey's multiple-comparison test; exact *p* values are shown in the panel.

**Supplementary Fig. 5. Individual tumor responses and immune memory after tumor rechallenge.**

**(a)** Individual tumor growth curves after NR, S-PRT, or F-PRT in combination with immunotherapy. Each curve represents one mouse. NR (n=10), S-PRT (n=10), and F-PRT (n=10) are shown in blue, red, and green, respectively.

**(b)** Rechallenge experimental design. Mice with complete tumor response and no subsequent late relapse after S-PRT plus immunotherapy or F-PRT plus immunotherapy were rechallenged with $2 \times 10^5$ EO771 cells injected into the contralateral right mammary fat pad. In the absence of tumor recurrence 15 days after rechallenge, mice were sacrificed for immune profiling of the spleen and left axillary tumor-draining lymph nodes by flow cytometry.

**(c-d)** Flow cytometry analysis of memory T-cell populations in the left axillary tumor-draining lymph nodes (c) and spleen (d) after tumor rechallenge. Groups included Control NR mice (n = 7) and mice previously treated with S-PRT + immunotherapy (n = 7) or F-PRT + immunotherapy (n = 8) that achieved a complete tumor response and remained relapse-free before rechallenge. CD4+ and CD8+ central-memory and effector-memory T-cell populations were analyzed. Boxes show the median and interquartile range, whiskers indicate minimum and maximum values, and each point represents one mouse. Comparisons were performed using ordinary one-way ANOVA.

**Supplementary Fig. 6. Flow-cytometry gating strategies.**

Representative gating strategies used for **(a)** T-cell immune profiling, **(b)** identification of myeloid-derived suppressor-cell populations, **(c)** macrophage phenotyping, **(d)** identification of memory T-cell populations, and **(e)** assessment of T-cell proliferation and activation. Gating was performed sequentially on singlets, viable cells, and $CD45^+$ leukocytes before identification of the indicated immune-cell populations.
