## Supplementary Table 1 for "Proton FLASH radiotherapy enhances control of triple-negative breast cancer through STING–IRF3 and CD8+ T-cell immunity"

**Supplementary Table 1. Flow cytometry antibodies, fluorochromes and clones.**

| Marker / Antibody | Fluorochrome | Company | Catalog Number | LOT Number | Clone |
|---|---|---|---|---|---|
| Live/Dead | Fixable Aqua Dead Cell Stain Kit | Thermo Fisher Scientific | L34965 | 3417064 | |
| CD45 | Pacific Blue | BioLegend | 157212 | B479797 | S18009F |
| CD3 | APC | eBioscience | 17-0032-82 | 2634921 | 17A2 |
| CD4 | PerCP/Cy5.5 | BioLegend | 100434 | B477454 | GK1.5 |
| CD8 | PE/Cy7 | eBioscience | 25-0081-82 | 2891539 | 53-6.7 |
| CD25 | Alexa Fluor 488 | eBioscience | 53-0251-82 | 2689384 | PC61.5 |
| FOXP3 | PE | eBioscience | 12-4771-82 | 2819238 | NRRF-30 |
| CD45 | PerCP | BioLegend | 103130 | B49134 | 30-F11 |
| CD11b | FITC | BioLegend | 101206 | B286843 | M1/70 |
| F4/80 | APC | BioLegend | 123116 | B298926 | BM8 |
| CD206 | eFluor 450 | eBioscience | 48-2061-82 | 2689355 | MR6F3 |
| CD80 | PE/Cy7 | BioLegend | 104734 | B413526 | 16-10A1 |
| Arginase 1 | PE | eBioscience | 12-3697-82 | 2699812 | A1exF5 |
| Ly6G | Pacific Blue | BioLegend | 127612 | B316649 | 1A8 |
| Ly6C | PE | eBioscience | 12-5932-82 | 2844137 | HK1.4 |
| Gr-1 | APC/Cy7 | BioLegend | 108424 | B327136 | RB6-8C5 |
| CD11c | APC | BioLegend | 117310 | B331091 | N418 |
| CD3 | eFluor 450 | eBioscience | 48-0032-80 | 2131781 | 17A2 |
| CD8 | PE | BioLegend | 100708 | B289299 | 53-6.7 |
| CD4 | FITC | BioLegend | 100406 | b423787 | GK1.5 |
| IFNγ | APC | BioLegend | 505810 | B396246 | XMG1.2 |
| Granzyme B | PerCP/Cy5.5 | BioLegend | 372211 | B357973 | QA16A02 |
| C3 | APC/Cy7 | BioLegend | 100222 | B292332 | 17A2 |
| CD44 | APC | BioLegend | 103012 | B472799 | IM7 |
| CD62L | Pacific Blue | BioLegend | 161208 | B458075 | W18021D |
| Ki-67 | PE/Cy7 | eBioscience | 25-5698-82 | 2335772 | SolA15 |
